# Pan-genome inversion index reveals evolutionary insights into the subpopulation structure of Asian rice (*Oryza sativa*)

**DOI:** 10.1101/2022.06.11.495682

**Authors:** Yong Zhou, Zhichao Yu, Dmytro Chebotarov, Kapeel Chougule, Zhenyuan Lu, Luis F. Rivera, Nagarajan Kathiresan, Noor Al-Bader, Nahed Mohammed, Aseel Alsantely, Saule Mussurova, João Santos, Manjula Thimma, Maxim Troukhan, Alice Fornasiero, Carl D. Green, Dario Copetti, Dave Kudrna, Victor Llaca, Mathias Lorieux, Andrea Zuccolo, Doreen Ware, Kenneth McNally, Jianwei Zhang, Rod A. Wing

## Abstract

Understanding and exploiting genetic diversity is a key factor for the productive and stable production of rice. Utilizing 16 high-quality genomes that represent the subpopulation structure of Asian rice (*O. sativa*), plus the genomes of two close relatives (*O. rufipogon* and *O. punctata*), we built a pan-genome inversion index of 1,054 non-redundant inversions that span an average of ∼ 14% of the *O. sativa* cv. Nipponbare reference genome sequence. Using this index we estimated an inversion rate of 1,100 inversions per million years in Asian rice, which is 37 to 73 times higher than previously estimated for plants. Detailed analyses of these inversions showed evidence of their effects on gene regulation, recombination rate, linkage disequilibrium and agronomic trait performance. Our study uncovers the prevalence and scale of large inversions (≥ 100 kb) across the pan-genome of Asian rice, and hints at their largely unexplored role in functional biology and crop performance.

## Main

Asian rice (*Oryza sativa*) is a staple cereal crop that has played an essential role in feeding much of the world for millennia^1,2^. As the population expands to almost 10-billion by 2064^3^, the rice community is searching for novel ways to breed new varieties that are sustainable, nutritious and climate resilient^2^. One source of the raw material required to meet this urgent demand is standing natural structure variation (SV), *i*.*e*. single nucleotide polymorphisms [SNPs], insertions/deletions [INSs/DELs], translocations [TRAs], and inversions [INVs] in the genomes of the more than 500,000 accessions of rice and its wild relatives that have been deposited in germplasm banks around the world^2^.

Inversions are an important subset of this natural variation tool box^4-6^ and have been shown to play important roles in genetic recombination (*e*.*g. Drosophila*^7,8^, *Helianthus*^9^, yeast^10^, bacterial^11^), genome evolution (*e*.*g*. mouse^12^, human^13,14^), and speciation (*e*.*g. Mimulus guttatus*^15^, chimps and humans^16,17^). In rice, inversions are understudied and have been limited to small and mid-size inversions as a consequence of the reliance on short-read data for their detection. For example, Wang et al. (2018) performed a genome scan of inversions in *O. sativa* cv. Nipponbare (*i*.*e*. IRGSP RefSeq) using re-sequencing data from 453 high-coverage genomes (> 20x) from the 3K Rice Genome Project (3K-RGP) and detected 152 ± 62 inversions per genome with a size range of 127.1 ± 19.4 kb^18,19^. A phylogenetic analysis of this dataset, including other SV data, demonstrated that SVs can be used to define the population structure of Asian rice^18^. Fuentes et al. (2019) went onto interrogate the entire 3K-RGP dataset in a similar manner and identified 1,255,033 inversions, with the vast majority falling in a size range of 50 bp – 500 kb^20^. A genome scan of the IRGSP RefSeq, plus reciprocal genome alignment to nine Asian rice and two AA-genome wild relatives (*i*.*e. O. rufipogon* and *O. longistaminata*) confirmed the presence a previously detected ∼5 megabases (Mb) inversion spanning the centromere of chromosome 6 in four *Xian*-*indica* (*XI*) varieties, relative to four *Geng*-*japonica* (*GJ*) varieties, as well as the two outgroup species^21^. A broad phylogenetic study in 13 cultivated and wild *Oryza* genomes using SVs resulted in the identification of 12 large inversions (*i*.*e*. 60-300 kb) that the authors inferred potentially led to the rapid diversification of the AA genome species within a 2.5 million years (MY) span^22^.

Although these studies contributed to a preliminary understanding of inversions in rice, they are limited due to their reliance on short-read sequencing technology, and the number and quality of genomes analyzed. Of note, a comprehensive analysis of inversions that utilizes ultra-high-quality reference genome sequences, which takes into account the population structure of Asian rice, remains uncharted. To reveal a comprehensive understanding of large inversions (≥ 100 bp) and explore their evolutionary impacts in Asian rice, we used a set of 15 platinum standard reference sequences (PSRefSeqs) that were sequenced with long-read sequencing technology, and assembled, edited and validated with a uniform pipeline^23,24^. When combined with the *O. sativa* IRGSP RefSeq^25^, these data can be used as a “pan-genome” proxy to represent the subpopulation structure of cultivated Asian rice, *i*.*e*. 15-subpopulations from subgroups *Geng/Japonica* (*GJ*), *Xian/Indica* (*XI*), *circum-Aus* (*cA*), *circum-Basmati* (*cB*), plus the largest admixed subpopulation, where K =15^24^. This Asian rice pan-genome was then scanned for inversions ≥ 100 bp, all anchored within a phylogenetic context, using two additional *de novo* assembled (to a similar quality) genomes from a representative species of the progenitor of Asian rice (*O. rufipogon*) and the BB genome species -*O. punctata*, as outgroups.

In this study, we comprehensively interrogated this pan-genome dataset to detect and analyze the inversion landscape of Asian rice at the population structure level, the results of which revealed salient evolutionary insights into the genome biology of Asian rice:

1. We created a novel Asian rice pan-genome, including 16 PSRefSeqs that represent its K=15 population structure, plus PSRefSeqs from two close wild relatives (*O. rufipogon* and *O. punctata*).
2. A pan-genome inversion index of 1,054 non-redundant inversions was generated and independently validated with physical maps (*i*.*e*. Bionano optical maps) and resequencing data (*i*.*e*. 3K-RGP).
3. A novel “pan-genome inversion rate” was estimated at 1,100 inversions per million years in Asian rice, which is 37 to 73 times higher than previous estimated in plants.
4. Biological functions *via* gene disruption, recombination rate, and linkage disequilibrium (LD) were investigated where we found that, on average, 8 genes were disrupted per genome; the genome recombination rate of a RIL population decreased from 6.98 to 4.00 cM/Mb; and 88.6% of the inversions tested may contain traces of recombination.
5. The biological consequences of a ∼400 kb inversion cluster (*i*.*e*. INV030400/INV030410/INV030420) on chromosome 3, that arose in the *Xian*/*Indica* (*XI*) and *circum-Aus* (*cA*) subgroups, was shown to be under positive selection, and was associated with delayed flowering with respect to standard genotypes.

## Results

### The 18-genome Data Package

To investigate the inversion landscape of Asian rice from a population structure perspective, we first combined a set of 16 previously published high-quality genomes that represent the K=15 population structure of *O. sativa*, plus the largest *Xian*/*indica* (*XI*) admixed subpopulation (*XI*-adm: Minghui 63 (MH63)) to create a “pan-genome” of Asian rice. To anchor this novel pan-genome within a phylogenetic context, we long-read sequenced, *de novo* assembled and validated two additional genomes from both a representative species of the progenitor of Asian rice - *i*.*e. O. rufipogon* [AA], and the African BB genome outgroup species - *O. punctata* (Table 1, Supplementary Table 1, Extended Data Fig. 1, and Supplementary Note 1).

All Asian rice assemblies were annotated using a uniform annotation pipeline to minimize methodological artifacts (Table 1, Supplementary Tables 2-4, Extended Data Fig. 2, and Supplementary Note 1), except for the *XI*-adm: MH63 and *XI*-1A: Zhenshan 97 (ZS97) genomes, whose annotations were previously published^23^. Lastly, we integrated and compared all annotations with that of the *GJ*-temp: IRGSP RefSeq^25^.

All 18 genomes, and their annotations, are henceforth referred to as the “18-genome data package” (See “18-genome data package” in the Supplementary Note 1 section for a complete description of this data set).

### Creation of a Pan-genome Inversion Index for Asian Rice

We pairwise compared 17 reference genome assemblies with the IRGSP RefSeq^25^ and identified a total of 2,915 inversions (≥ 100 bp) (Table 2, Supplementary note 2), of which, 1,054 were non-redundant (Supplementary Table 5). As expected, more inversions were observed when we compared the Asian rice pan-genome with both the AA and BB genome outgroups to the IRGSP RefSeq: 194 (total length = 13.05 Mb) and 316 (total length = 17.85 Mb) for *O. rufipogon* and *O. punctata*, respectively (Table 2). On average, each *O. sativa* genome was found to contain 160 inversions, ranging from 88 (*GJ*-subtrp: CHAO MEO) to 187 (*XI*-3B1: KHAO YAI GUANG) (Table 2). We found a larger number of inversions (172 to 187) when comparing the *O. sativa XI*-subgroup genomes to *GJ*-temp: IRGSP RefSeq, than when comparing the *O. sativa GJ*-subgroup genomes (88 to 112 inversions) to the same reference (Table 2). The total length of the inverted regions, per genome, ranged from 7.73 Mb (*GJ*-subtrp: CHAO MEO) to 14.95 Mb (*cA*2: NATEL BORO) (Table 2). When chromosome location was taken into account, these inversions appeared to be evenly distributed genome-wide (Kolmogorov-Smirnov test, *P* value 0.02-0.95) (Supplementary Table 6, Extended Data Fig. 3, Supplementary note 3).

### Species and Subpopulation Specific Inversions

Of the 1,054 non-redundant inversions detected (Supplementary table 5), we classified them into different categories: *i*.*e*. species-specific (the inversion could be only observed in *O. punctata, O. rufipogon* or *O. sativa* genomes); group-specific (only observed either in *GJ, XI, cA* or *cB* subgroup in *O. sativa*); and genome-specific (only observed in one of 16 *O. sativa* genomes). As a result, 968 (91.8%) appeared to be species-specific (*i*.*e. O. sativa*: 550 (totaling 50.21 Mb), *O. rufipogon*: 105 (11.05 Mb), and *O. punctata*: 313 (17.84 Mb)) (Fig. 1). The remaining 86 were found in two or more species and totaled to about 2.06 Mb in size.

**Fig. 1.**
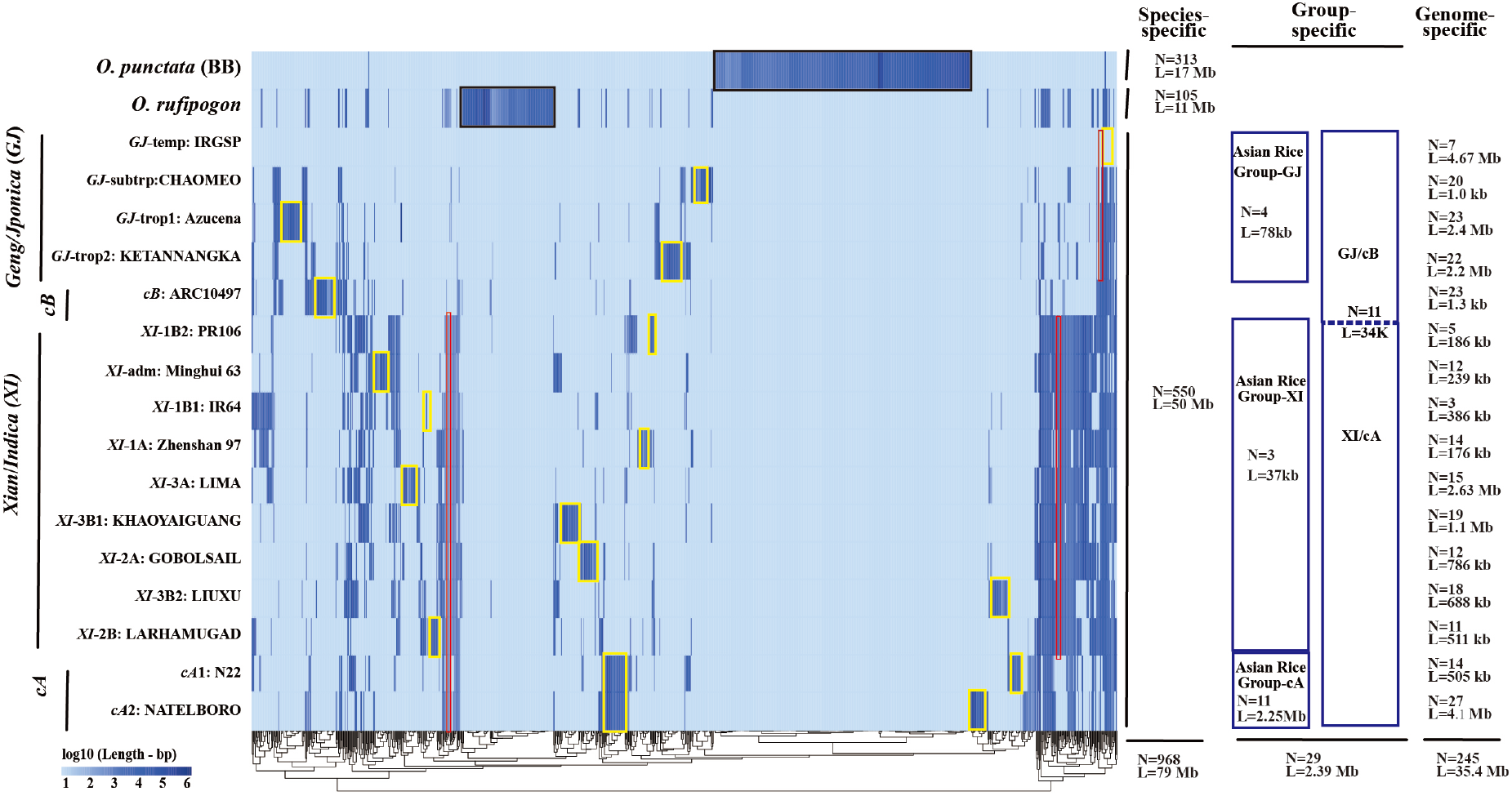
Inversion landscape across 17 PSRefSeqs all relative to the IRGSP RefSeq shows species-specific, group-specific and genome-specific inversions, *i*.*e*., species-specific inversions (*O. punctata* and *O. rufipogon*) are shown by black rectangles, group-specific inversions are shown by red rectangles, and genome-specific inversions are shown by yellow rectangles, respectively. On the left, accessions are phylogenetically ordered^24^, on the bottom, the tree are the clustering of inversions, and on the right, the numbers and lengths of the specific inversions are presented.

Two hundred and forty-five of the 550 *O. sativa* specific inversions were specific to one of the 16 *O. sativa* genomes, while 305 were shared with more than one genome (Fig. 1). The frequency of 100 randomly selected *O. sativa* genome-specific inversions were further investigated in different subpopulations using a subset of 3K-RGP data set (*i*.*e*. 192 highly re-sequenced (> 20 ×) accessions) (Supplementary table 7). With the exception of 18% of the inversions, for lack of evidence within the 3K-RGP dataset, the remaining 82% could be classified in four groups: 15% genome-specific inversions, 23% subpopulation specific inversions, 33% near-subpopulation specific, and 11% subpopulation shared inversions (See supplementary online methods for category definitions) (Supplementary table 8).

Of the 305 inversions present in more than one of the 16 Asian rice genomes included in our dataset, we identified 29 that were shared among closely related populations (Fig. 1), which we defined as “group specific”. Four inversions were shared among 4 *GJ* genomes, 3 were shared among 9 *XI* genomes, 11 were shared among 2 *cA* genomes, and 11 were identified by comparing 5 *GJ* and *cB* genomes to 11 *XI* and c*A* genomes (Fig. 1). These 29 inversions were also studied in different subpopulations across a high-coverage subset of 3K-RGP dataset. Excluding two inversions that couldn’t be tested (*i*.*e*. because no reads were observed at the breakpoints), we found that 11 (38%), 13 (44%) and 3 (10%) inversions were group specific, near-group specific and group shared inversions at the subpopulation level, respectively (Supplementary table 8). The remaining 278 inversions appear to be shared across different genomes or subpopulations reflecting the substantial admixture in the evolution of subpopulations in Asian rice and mixed ancestry in the pedigrees of some accessions used (*e*.*g*. IR8, IR64, MH63, and ZS97)^26,27^.

Altogether, > 85% of the genome specific inversions and > 85% of the *O. sativa* group specific inversions could be validated with the high-coverage subset of the 3K-RGP dataset, and appear to be Asian rice subpopulation(s)-or subgroup(s)-specific. This analysis validates the accuracy in detecting inversion boundaries, provides initial estimates of inversion frequencies in rice subpopulations, and patterns of shared inversions between subpopulations.

### Five Largest Inversions

We identified 5 inversions greater than 1 Mb relative to the IRGSP RefSeq on chromosomes 1, 6, 8, and 10 (INV010130 [2.00 Mb], INV010560 [1.82 Mb], INV060390 [4.57 Mb], INV080710 [1.12 Mb] and INV100690 [1.33 Mb]), two of which (INV060390^28^ and INV080710^29^) were previously reported.

INV010130, INV010560 and INV060390 appear to be Asian rice specific (Fig. 2, Supplementary Table 5 & 9, Supplementary note 4), and could not be found in the outgroup genomes. To determine if these inversions are subpopulation(s) specific, we interrogated the high-coverage subset of the 3K-RGP data set and found that INV010130 was specific to the *XI*-3A subpopulation, INV010560 to the c*A* and *XI*-adm subpopulations, and INV060390 to the *GJ*-tmp and *GJ*-subtrp subpopulations (Supplementary Table 9, Extended Data Fig. 4, Supplementary note 4).

**Fig. 2.**
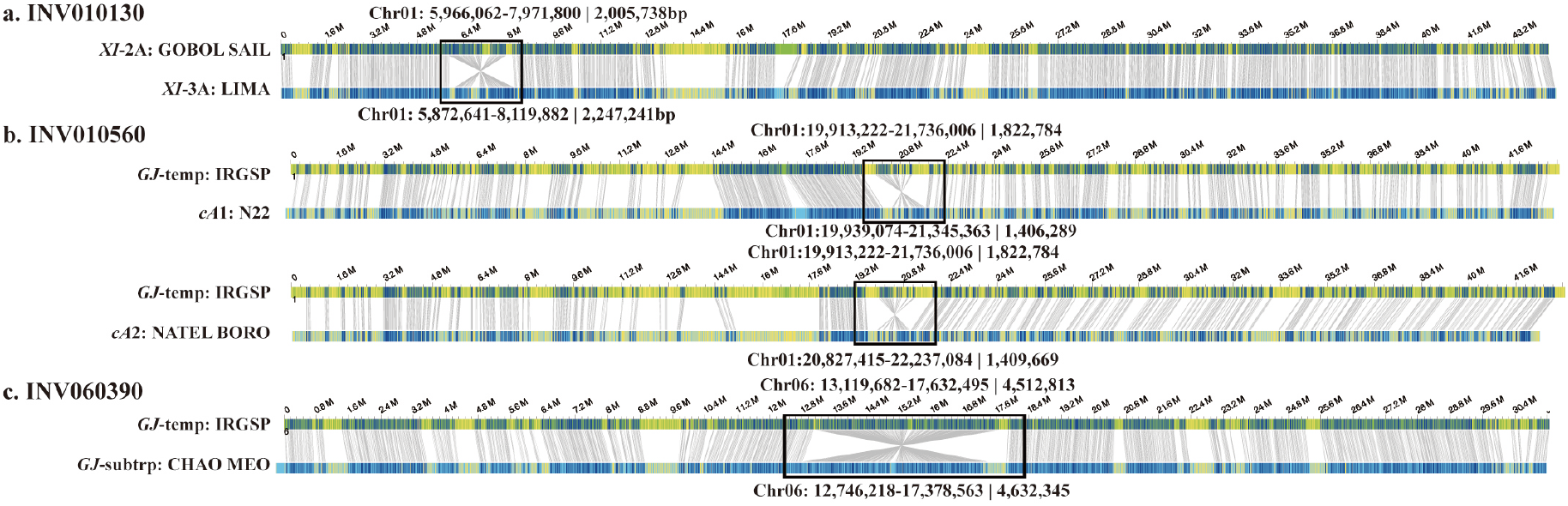
Bionano validation of inversions larger than 1Mb. a. INV010130, b. INV010560 and c. INV060390. In each panel, the top line has the optical map used as a reference, the bottom line has the genome assembly of the variety with the inversion. Gray lines connect restriction sites that are aligned (blue regions), while yellow segments are unaligned regions. Black boxes highlight the position of the inversion.

The remaining two inversions (*i*.*e*. INV080710 and INV100690) were detected only in *O. punctata* and *O. rufipogon*, respectively (Supplementary Table 5 & 9), and thus appeared to be species specific. To test this hypothesis, we investigated the presence or absence of these inversions in high-quality genomes of 5 additional *Oryza* species (*i*.*e. O. nivara* [AA], *O. glaberrima* [AA], *O. barthii* [AA], and the distantly related subgenomes of *O. coarctata* [KKLL] and *O. alta* [CCDD] (unpublished data). Results showed that neither of these inversions could be detected in these five species. Thus, we conclude that INV080710 (*O. punctata*) and INV100690 (*O. rufipogon*) are species specific.

### Characterization of Transposable Element Content within Inversions and Breakpoints

Transposable elements (TEs) are known to be associated with inversions^18,20,30^, thus we analyzed the TE content across the inversion index, and at their breakpoints. The total amount of TE related sequences within these inversions ranged from 64% (*GJ*-subtrp: CHAO MEO) to 73% (*XI*-adm: MH63) (Supplementary Table 10), which is significantly (student’s test, *p* < 0.01) higher than the average content of TEs across all 16 *O. sativa* genomes at 51.3% (Table 1, Supplementary table 10 & 11). These results demonstrate that TEs are enriched within inversions.

Analysis of breakpoints revealed that both long terminal repeat retrotransposons (LTR-RTs, *i*.*e*. Ty3-*gypsy* and Ty1-*copia*) and DNA TE Mutator-like elements (MULEs) were significantly (student’s test, *p* < 0.01) enriched, when the frequency of their presence at the 2,108 breakpoints was compared to 21,080 randomly selected genomic locations (*i*.*e*. 10 replicates) (Fig. 3a). We further studied TEs at the breakpoint of each inversion that were shared across all Asian rice genomes. In doing so, we identified 17 TE families (*i*.*e*. 13 Ty3-*Gypsy*, 1 Ty1-*Copia*, 2 *CACTA*, and 1 *Mutator*) present at the breakpoints of more than 10 inversions (Fig. 3b & Supplementary Table 12). An example of an inversion enriched in TEs, including the internal and LTR portions of at least three different LTR-RTs, is shown in Fig. 3c.

**Fig. 3.**
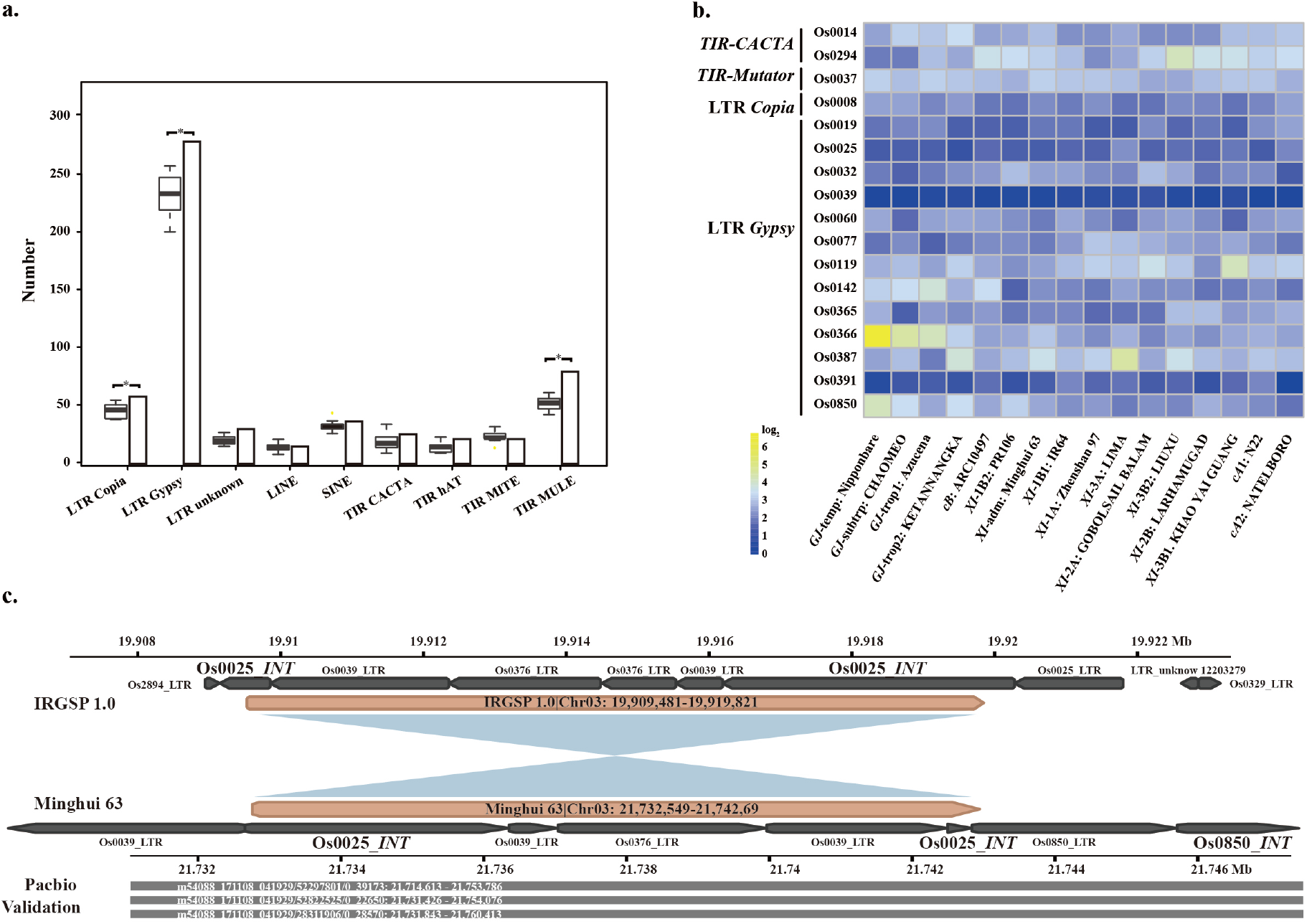
Transposable elements (TEs) are associated with inversions. a. The amount (y-axes) of different TE families (x-axis) show that three TE families (*i*.*e*. LTR-RT Ty1-*copia*, Ty3-*gypsy* and DNA-TE MULE) were observed in higher frequencies at the breakpoints of the pan-genome inversion index than the resampled control tests. Box-plots and bar-plots show the frequencies of TEs observed at the breakpoints of 10 random resampling regions and the pan-genome inversion index, respectively. b. Enrichment/depletion of 17 TEs present at the inversion breakpoints with more than 10 copies. c. Details of Ty3*-gypsy* Os0025 presence at inversion breakpoints, with support from CCS PacBio long reads.

Together, our results reveal an enrichment of TE related sequences both within inversions and at their breakpoints.

### Characterization of Gene Content within Inversions and Breakpoints

Based on the pan-genome inversion index we identified a total of 15,530 genes (∼1,035/genome) within or at inversion breakpoints (Supplementary Table 13). To investigate the effect of inversions on the expression of genes located within inverted regions, we interrogated a transcriptome dataset derived from a subset of the 18-genome data packaged including *O. sativa cv. XI*-adm: MH63, *XI*-1A: ZS97 and *GJ*-temp: Nipponbare (*i*.*e*. dataset#2 -see online methods). Based on 284 and 356 expressed orthologous genes between the reference (*GJ*-temp: IRGSP RefSeq) and two queries (*XI*-adm: MH63 and *XI*-1A: ZS97), we detected 10.9% (31) genes from *XI*-adm: MH63 and 7.3% (26) from *XI*-1A: ZS97 that were differentially expressed (DEG, fold change > 2, *P* value < 0.01) (Supplementary table 14) relative to the *O. sativa cv. GJ*-temp: Nipponbare genome.

To investigate the effect of inversions on the transcription of genes located at inversion breakpoints -*i*.*e*. about 55 genes per genome (Supplementary table 13), we interrogated both our baseline RNA-Seq dataset (dataset#1-see online methods) and dataset#2 for changes in transcript abundance. On average, 28 of the 55 genes per genome were found to be expressed in the tissues tested (Supplementary table 14). Of these, transcript abundance of an average of 20 genes per genome did not change due to the presence of duplicated genes at both ends of their inversion breakpoints (Supplementary table 14). An example of this observation is represented by two *OsNAS* genes (*NAS1* and *NAS2*) located at the breakpoint of INV030200 (∼4.3 kb) (Fig. 4a & a). The remaining ∼8 genes/genome were single copy and were disrupted by the inversion events, leading to the absence of transcript evidence (Supplementary table 14). As an example, transcripts of the Nipponbare Fbox gene (Os11g0532600) can be detected in the four tissues tested. However, the first exon of this gene is disrupted in MH63 by INV110960, resulting in transcript ablation (Fig. 4C & D).

**Fig. 4.**
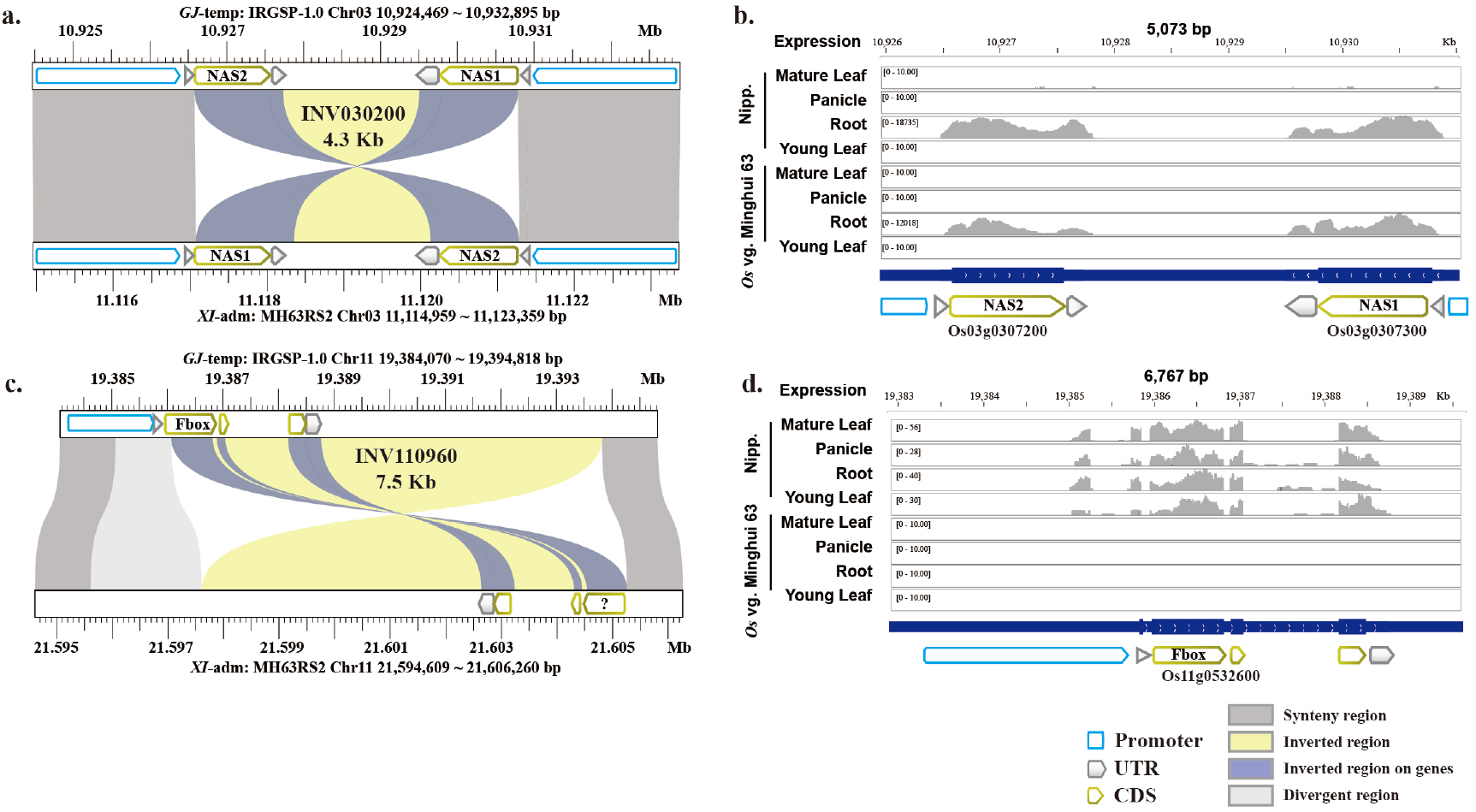
Transcript abundance of genes located at inversion breakpoints. a. Two copies of the *OsNAS* gene lie at the end of an inversion in the MH63RS2 (*XI*-adm) genome. This inversion disrupted the 5’ UTR regions. b. *OsNAS* gene transcript abundance in root tissue. c. The coding sequence (CDS) of a Fbox gene was disrupted by an inversion in the MH63RS2 (*XI*-adm) genome. d. Fbox transcript abundance was suppressed in all tissues tested in the MH63RS2 (*XI*-adm) genome.

### Recombination Rate and Genomic Inversions

To evaluate the effect of inversions on recombination frequency, a previously published recombinant inbred line (RIL-10) population of 210 inbred lines^31^ derived from a cross between *O. sativa cv. XI*-adm: MH63 and *XI*-1A: ZS97 was investigated. We detected 78 inversions between MH63 and ZS97, totaling 3.58 Mb and 3.51 Mb based on the MH63RS2 and ZS97RS2 genome assemblies, respectively (Supplementary table 15). The recombination rate along each chromosome was assessed by comparing genetic and physical distances between neighboring bins. The average recombination rate for each chromosome ranged from 5.95 (chromosome 6) to 9.92 (chromosome 12) cM/Mb, and varied from 0 to 153.93 cM/Mb across the genome with an average of 6.98 cM/Mb (Extended Data Fig. 5A). The average recombination rate over the 78 inverted regions was 4.00 cM/Mb (0 - 23.26 cM/Mb), which is significantly lower (Student’s t-test, *p* = 0.0002) than that observed genome-wide (Extended Data Fig. 5B). These results indicate that a marked suppression of genetic recombination is associated with inversions.

### Effect of Large Inversions on Population SNP Variation

The occurrence of inversions can affect DNA polymorphism at the population level in several ways, including increased divergence in the inverted region and changes in linkage disequilibrium (LD) patterns^32^. The latter is particularly interesting as it can affect SNPs that are mapped to positions megabases apart, and can be a confounding factor in LD-based analyses. To determine whether large *O. sativa* inversions left a trace in patterns of LD along the IRGSP RefSeq, we used the 3K-RGP dataset to examine LD blocks near inverted regions (> 100 kb). First, inversions having a reciprocal overlap of more than 80% of their length were clustered and considered as putative unique inversions. In doing so, we considered 53 clusters including from 1 to 6 inversions each (Supplementary Table 16). An inversion fixed in a population may lead to the disruption of LD blocks, in which some SNPs flanking the inversion on one side are in LD with SNPs on the distal part of the inversion, but not on the adjacent part (Fig. 5), due to the reversed order of SNPs inside the inverted region in samples that carry the inversion allele. By an LD block we mean only a set of SNPs in high LD (r^2^ > 0.8 in this analysis).

**Fig. 5.**
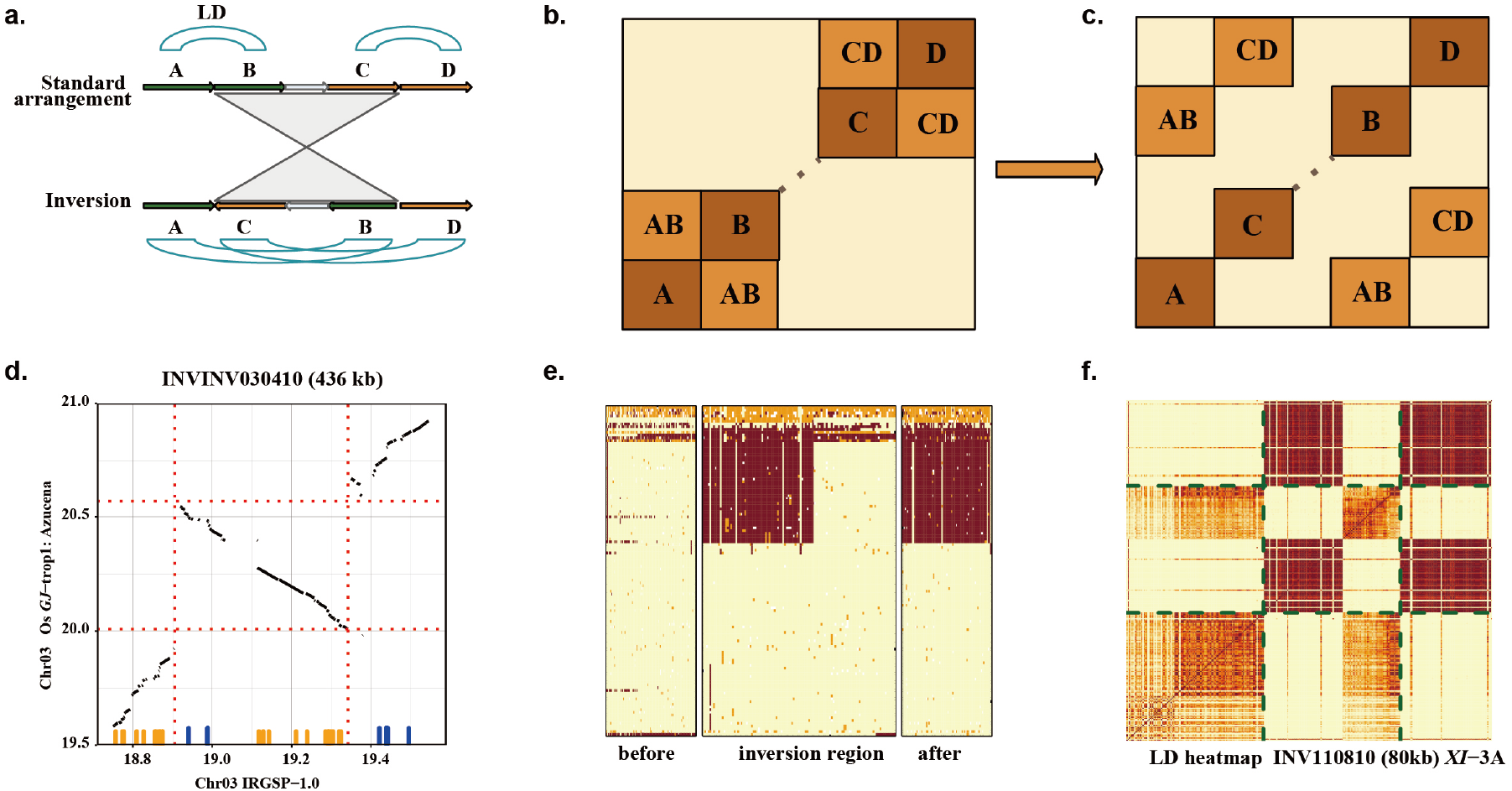
Population level SNP variation across large inversions. A schematic diagram of LD block disruption arising from the presence of an inversion, as shown in A and B. a. Cartoon view of an inversion with breakpoints disrupting two LD blocks. b. Expected features of the corresponding LD heat map. c. Example of SNP blocks in high LD that are disrupted by an inversion. d. The panel shows alignments, with the inversion marked by dotted lines. Small vertical lines above the horizontal axis mark the location of SNPs constituting a disrupted LD block. Orange and blue colors delineate two LD blocks that are contiguous in the of *GJ*-trop1 population, but appear as split when aligned to the IRGSP RefSeq (*GJ*-temp). Disruption of Azucena (*GJ*-trop1) haplotype blocks along the IRGSP RefSeq in the region of INV030410, as shown in e and f. e. Genotype heat map of the *GJ*-trop1 subpopulation (samples in rows, SNPs in columns; light yellow: reference call, orange: heterozygous, brown: homozygous variant). f. LD heat map of the same subpopulation. Dotted lines show the inversion region. Darker colors show larger r^2^. Note that the scaling of X-axis in the genotype heat map is not uniform, allotting half of X axis space to the inverted region.

Next, we examined the entire 3K-RGP variation data set and searched for LD blocks that connect the flanking regions of inversions, having no SNPs in the proximal parts of each inversion. Such blocks (Fig. 5), were found in nearly all large inversions (63 out of 81 [75.3%] alignment-based inversions, or 47 out of 53 [88.7%] inversion clusters) (Supplementary Table 16) with only two classes of exceptions: *i*.*e*. inversions in regions of complex chromosomal rearrangements (INV080210-INV080250, INV080510-INV080530, INV110600-INV110660), and three putative “recent” inversions (INV020230, INV100080, INV100320), each of which were found in single genomes and may lack sufficient frequencies in a population to contain traces of recombination. Some of the disrupted LD blocks contained a particularly large number of SNPs and were seen as a distinctive checkered pattern on LD heat maps (Fig. 5). This comparatively large number of SNPs along with low haplotype diversity, despite the presence of recombination, could be a consequence of selective pressure.

### Phenotypic consequences of inversions: Inversion Cluster 92

To investigate the phenotypic consequences of inversions (if any) on agronomic traits in rice, we correlated ten known phenotypes catalogued in SNP-Seek (https://snp-seek.irri.org/) across the high-coverage subset of the 3K-RGP data set (*i*.*e*. same subset as mentioned above) with our pan-genome inversion index. In this analysis, we used a linear model function in R to assess association between phenotype and inversion status, controlling for population structure, and in doing so, we identified an inversion cluster (*i*.*e*. INVCluster92), 400 kb in length, which was significantly (linear model test, *p* < 0.01) associated with delayed flowering time of 13 days, on average (Fig. 6, Supplementary table 17).

**Fig. 6.**
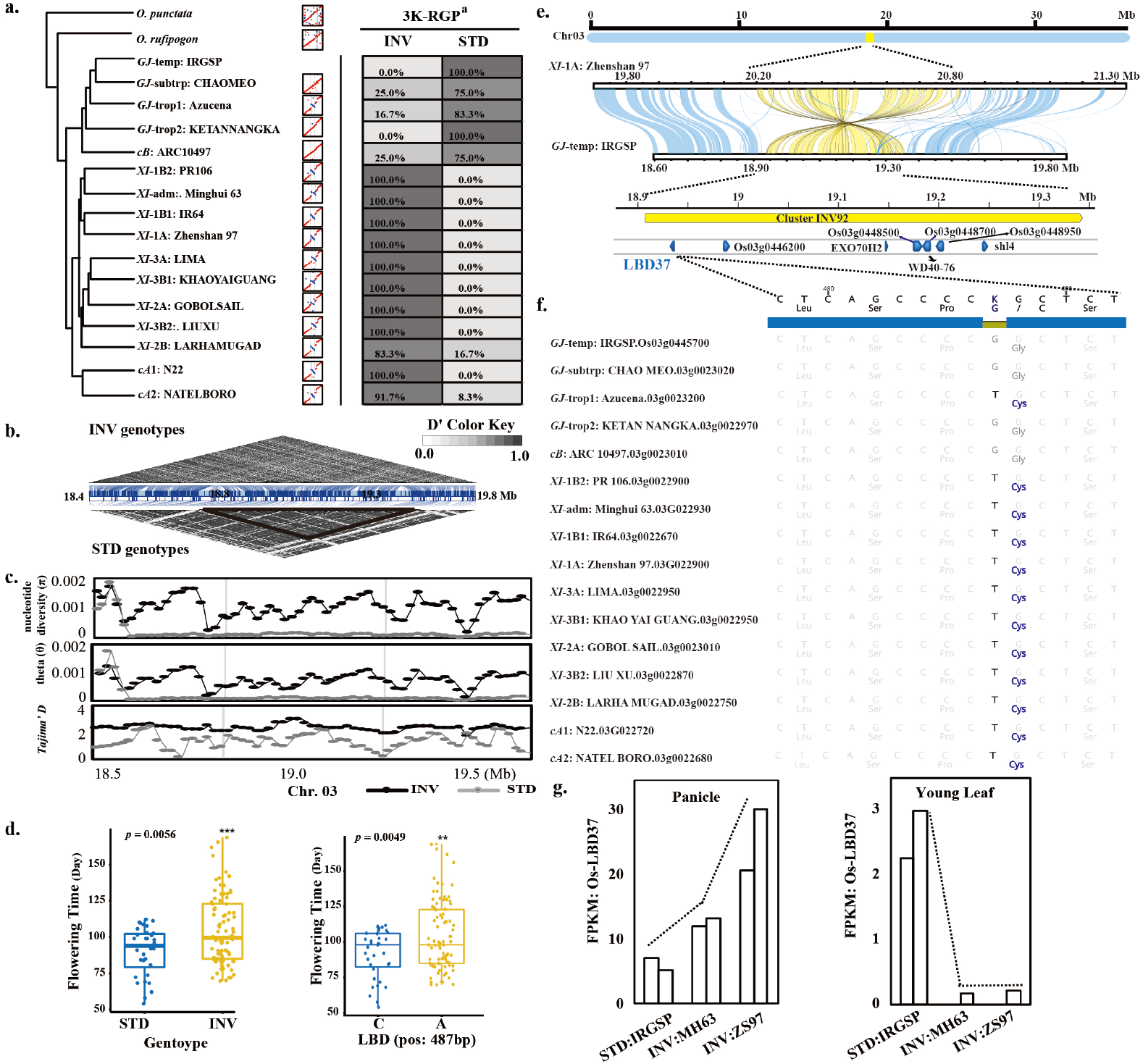
An integrated study of ClusterINV92. a. Genome comparisons identified that inversion ClusterINV92 arose in *XI* and *cA* genomes and are nearly fixed in *XI* and *cA* subpopulations. Note: ^a^ means 192 deep re-sequenced (> 20 ×) samples from the 3K-RGP were used for validation. b. LD block disruption arising from the presence of ClusterINV92in populations with INV genotypes compared to standard (STD) genotypes. c. Population genetic variation of nucleotide diversity, theta, and Tajima’s *D* of inversion cluster 92 genotypes compared to standard (STD) genotypes. The gray vertical lines delimit the inversion coordinates on the IRGSP reference. d. Phenotyping test shows a significant difference (linear model test, *p* < 0.01) in flowering time between ClusterINV92 genotypes and standard (STD) genotypes (left), and SNP variation in the LBD gene (right). e. A total of 8 genes, including 4 reported genes, were observed in ClusterINV92. f. A single SNP (G to T) caused a non-conservative amino acid change from the hydrophobic glycine (Gly) to hydrophilic cysteine (Cys) within the LBD gene. g. Expression of the LBD gene was up-regulated in panicle tissue, and suppressed in young leaf tissue of INV genotypes *XI*-adm: MH63 and *XI*-1A: ZS97, compared with the standard IRGSP RefSeq genotype.

INVCluster92 was found to be composed of three inversions (*i*.*e*. INV030400/INV030410/INV030420), with a minor difference at the breakpoints (Supplementary table 5), and is shared among twelve of the sixteen genomes in our Asian rice pan-genome data set (*i*.*e*. 9 *XI*, 2 *cA* and 1 *GJ* genomes (*GJ*-trop1: Azucena)) (Fig. 6a). Analysis of the 3K-RGP subset revealed that 137 (71.4%) contained the inversion (INV) genotype, while 55 (28.6%) did not (*i*.*e*. standard genotype (STD)) (Fig. 6a). Comparative LD analysis of STD vs. INVCluster92 genotypes is shown in Fig. 6b, and showed that the LD block of STD genotypes was disrupted by INVCluster92. In addition, the INV genotypes showed a significantly higher (student’s test, *p* < 0.01) nucleotide diversity (π), Watterson’s theta (θ), and Tajima’s *D* than the STD genotypes (Fig. 6c). This evidence supports the hypothesis that the regions immediately surrounding INVCluster92 may be under positive directional selection.

To find clues as to why INVCluster92 is associated with delayed flowering time (Fig. 6d, left), we compared gene content, structure and expression differences for eight genes present in both the STD and INV genotypes. Of note, four of these genes (*i*.*e*. LBD37^33^, EXO70H2^34^, WD40-76^35^, shl4^36^) have been functionally characterized (Supplementary table 18), and 4 are unknown (Fig. 6e). Among these eight genes, a total of 3 SNPs and a 12-bp insertion were observed within the coding sequence (CDS) regions of two genes - LBD37 (Os03g0445700) and WD40-76 (Os03g0448600). For LBD37, a single ‘G’ to ‘T’ SNP (SNP-1) was observed at base pair 487 resulting into a non-conservative amino acid change from glycine to cysteine (Fig. 6f). For gene WD40-76, 4 ‘GCC’ repeats were inserted (in frame) 6 bp after the start codon, resulting in the addition of 4 alanine residues at the beginning of the protein. In addition, two additional SNPs were detected in this gene, a synonymous ‘A’ to ‘G’ SNP (SNP-2) at position 468 bp and an ‘A’ to ‘C’ SNP (SNP-3) at position 557bp that changed a charged histidine amino acid into a non-polar proline amino acid (Extended Data Fig. 6). We also validated these SNPs across the 3K-RGP high-coverage subset and found that all these natural variants were absent in all (55) STD genotypes (Supplementary table 19). However, SNP-1 and the 12 bp InDel were present in 97.8%, SNP-2 was found in 90.5%, and SNP3 was found in all (137) INV genotypes.

Preliminary transcript abundance analysis of the eight genes were compared using deep RNA-Seq data from leaves, roots and mixed stage panicles (*i*.*e*. RNA dataset#2). The only difference in transcript abundance that could be observed was for gene LBD37 in panicle and young leaves, respectively. The LBD37 gene appeared to be up-regulated in panicle tissue, but down-regulated in young leaf tissues in INV genotypes, as compare with STD genotypes (Fig. 6g and Supplementary table 18), which is compatible with the inversion genotype and SNP-1. The phenotype variation analysis based on SNP-1 is also congruent with LBD37 over-expression (Fig. 6d, right), as previously reported in rice, *i*.*e*. a delay in heading date^33^. These results suggest that SNP-1 within LBD37 is under positive selection and may contribute to the observed phenotypic variation.

## Discussion

Inversions are an important class of structural variations that have been shown to play important roles in the suppression of recombination that can lead to the selection of adaptive traits, reproductive isolation and eventual speciation, and are quite common in plants^22,32,37^. For example, over the 50-60 MY history of the *Poaceae*, where gene order has been largely conserved, Ahn and Tanksley (1993) showed (using molecular genetic maps) that multiple inversions and translocations occurred during the evolution of maize and rice from a common ancestor^38^.

Here, we present, to our knowledge, the first comprehensive analysis of the inversion landscape of any cereal at the population structure level with the discovery of 1,054 non-redundant inversions that range from 8 Mb to 25 Mb in cumulative size (Table 2). It is estimated the AA genomes of the *Oryza* diverged from the BB genome type about 2.5 million years ago (MYA)^2^, which equates to an inversion rate of 63.2 inversions per million years (*i*.*e*. 316 inversions/ (2*2.5 MY)) - about 2 to 4 times higher than that recently estimated in plants (*i*.*e*. 15 to 30 inversions/MY)^32^. However, Huang and Rieseber^32^ (2020) noted that this earlier estimate should be considered an underestimate and is dependent on the quality of the genomes analyzed, and other factors. If we use the implied AA genome diversification rate of ∼0.50 net new species/million years^2,22^, then we calculate an inversion rate of 194 *O. sativa* inversions/MY - about 6.5 to 13 times higher than recently estimated in plants^32^. Taken one step further, by taking into account that Asian rice is estimated to have been domesticated 10,000 years ago^2^, and using only divergence within GJ group genomes not shared by any other groups (22 out of 88 inversions in KETAN NANGKA) which is likely post domestication, we can arrive at an estimated inversion rate > 1,100 inversions/MY (*i*.*e*. > 37 to 73 times previous estimates in plants^32^). Such a rapid pan-genome inversion rate over such a short time period may be reflective of high fixation rates of rearrangements in plants^6^, high chromosomal evolution rates in annual plants^39-41^, and intense human selection since the dawn of agriculture^18,20,42^.

Although an inversion genome scan for rice has been previously published^20^, when we extracted the inversion coordinates (*i*.*e*. 2,402 inversions, average length 43.3 kb), from the same set of 15 accessions used to generate our pan-genome mapped to the IRGSP-RefSeq, we found that only 200 could be validated with dot plots (a 91.7% false positive rate) (Supplementary Table 20), 194 of which overlapped with our inversion index. The 6 remaining contained 2 that overlapped, and only 4 that were not present in our inversion index (Supplementary Table 21). These analyses combined reveal the limitations of inversion callers with short read data and provide a cautionary note as to the validity of many of the inversions catalogued to date.

Several key factors led to our ability to generate a definitive gene inversion index for cultivated Asian rice. The first was our use of a set of 16-ultra high-quality reference genomes that represented the K = 15 population structure of Asian rice^24^, and 2 phylogenetically anchored wild AA and BB genome species (Table 1 and Supplementary Table 1). Secondly, we did not computationally collapse this 18-genome data package into pan-genome (*e*.*g*. genome graph), but maintained all 18 genomes in their native state. This was key to our ability to precisely compare all genomes one-by-one. Lastly, we interrogated a high-sequence coverage subset of the 3K-RGP data to estimate and validate the population genetics of each inversion. As sequencing costs continue to plummet, the ease at which ultra-high-quality genomes can be generated, and with computational power exceeding current limits^43-45^, we predicted that there will no longer be a need to computationally generate pan-genomes to perform similar analyses across much larger genomes, such as wheat (genome size = 15 Gb)^46^.

The Asian rice pan-genome inversion index is the first step on our quest to precisely discover all standing natural variation that exists in Asian rice and eventually the genus *Oryza* as a whole. The next step will be the generation of a digital genebank for Asian rice whereby resequencing data from > 100,000 accessions will be mapped to our pan-genome - *i*.*e*. all 16 genomes. Preliminary data (unpublished) shows that we can now easily call SNPs with resequencing data from > 3,000 individuals in 5 days per genome or less using supercomputer workflows optimized for GATK4^47^ software. Such call rates will undoubtedly increase over the next year with a targeted rice digital genebank release date of January 1^st^, 2025.

### Online Methods

All the methods are available in the supplementary information.

## ACKNOWLEGEMENTS

This research was supported by King Abdullah University of Science & Technology’s Baseline funding, and the University of Arizona’s Bud Antle Endowed Chair for Excellent in Agriculture to R.A.W., Huazhong Agricultural University’s Start-up Fund, and Fundamental Research Funds for the Central Universities (2662020SKPY010) to J.Z; USDA ARS 8062-21000-041-00D to DW. The authors acknowledge support from the Shaheen Cray XC40 platform at KAUST Supercomputing Laboratory, the computing platform of the National Key Laboratory of Crop Genetic Improvement at HZAU, Peter VanBuren for systems support and the Elzar High Performance Computing facility (NIH S10 OD0286321-01) at Cold Spring Harbor Laboratory for providing the computational resources, and help from the Persephone team that hosts our 18-genome data package on their portal for genome visualization.

## AUTHOR CONTRIBUTIONS

A.Z., D.W., K.M., J.Z., and R.A.W. designed and conceived the research. K.M. and IRRI provided seed and/or tissue for all *Oryza* accessions. D.K., N.M., D.Ch., and M.L. performed DNA extractions and genome sequencing. Y.Z., Z.Y., and J.Z. performed sequence assembly, GPM edit and validation of 18 genome sequences. Y.Z., N.M., D.K. and V.L. carried out the optical map sequence and analysis. D.Ch., Y.Z., J.S. and K.M. performed population genetic analysis. K.C., Z.L., and D.W. performed the genome annotation and validation. Y.Z., Z.Y., J.Z. and A.Z. performed the gene expression analysis and TE annotation. Y.Z., A.Z., J.S., A.A., S.M., Z.Y., and J.Z. carried out the inversion identification and population level validation. L.R., N.K., M.T., M.T., C.D., and K.C. managed the computing platforms. Y.Z., Z.Y., D.Co., K.C., N.A., A.F., A.Z., K.M., J.Z., and R.A.W wrote and edited the paper. All authors read and approved the final manuscript.

## Figs legends

## Extended Data Figs

**Extended Data Fig. 1.**
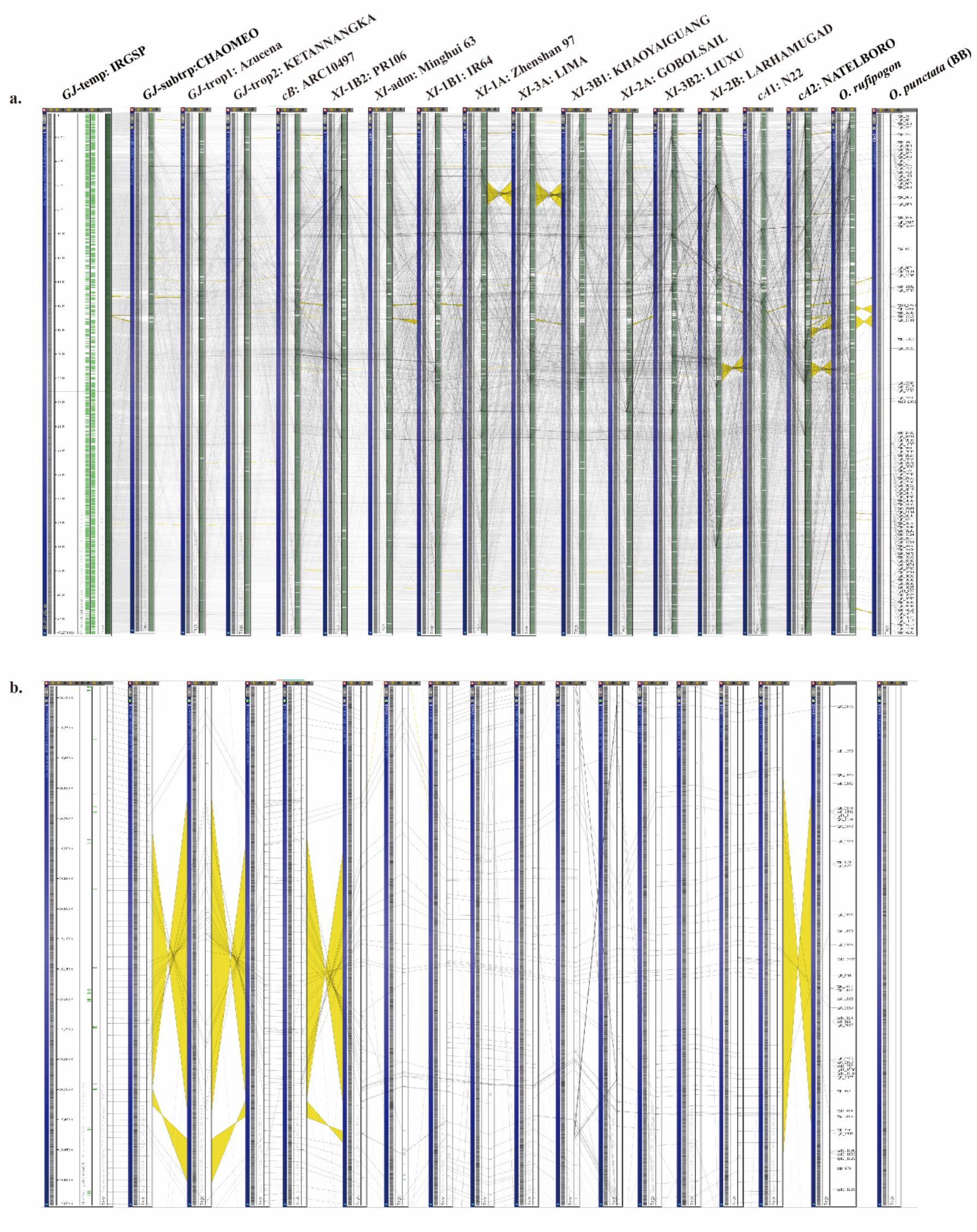
The 18-genome dataset (18 PSRefSeqs) was input into Persephone (https://web.persephonesoft.com/) and made publicly available. a. A panel shows overall alignments of the 18 maps (genomes) using chromosome 1 as an example. The gray lines show the alignments of sequence tags and the yellow ribbons show the inversions. b. A panel shows an 800 kb region that includes INVcluster92 with yellow ribbons.

**Extended Data Fig. 2.**
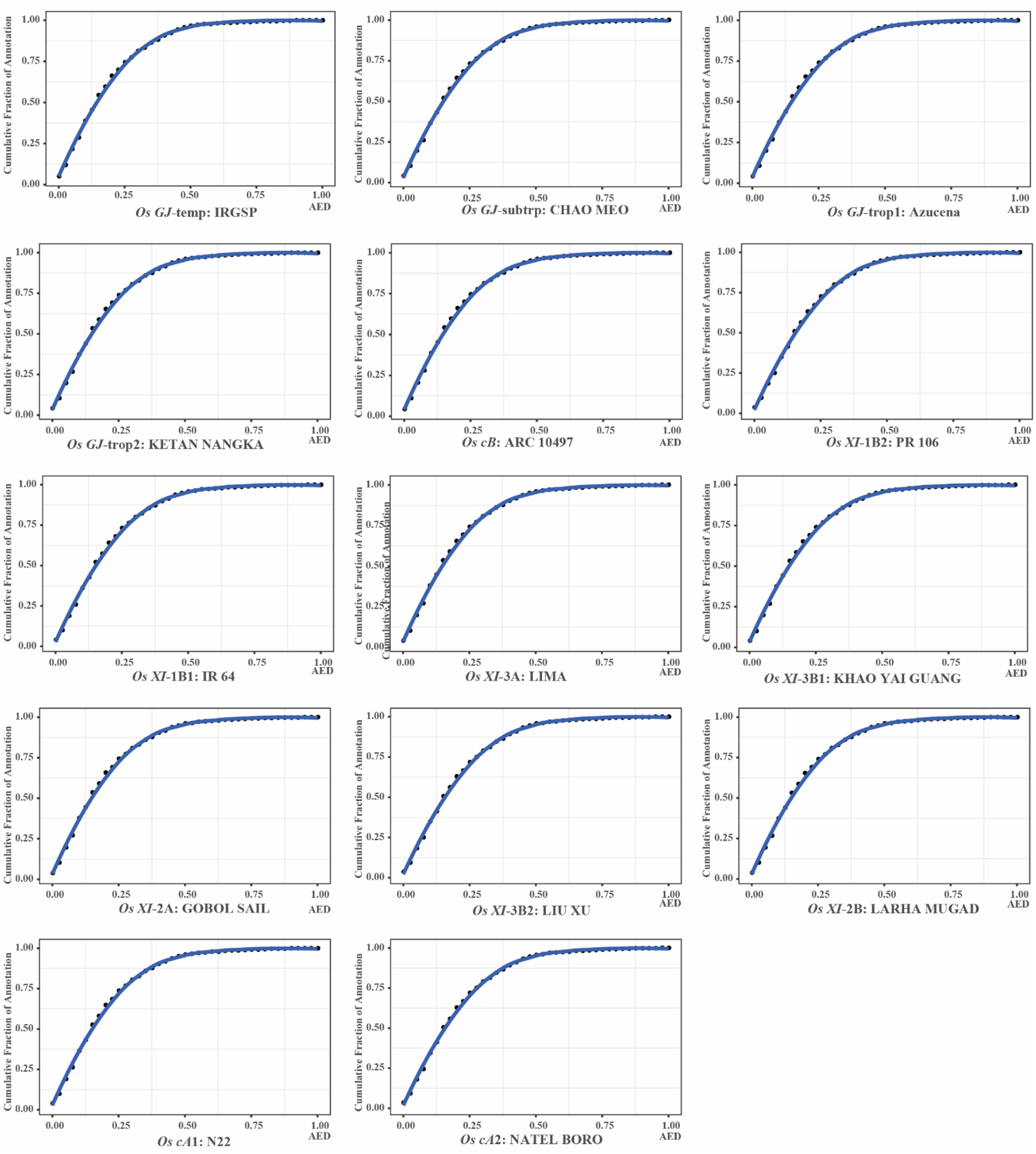
Cumulative AED distributions of 13 genomes and their annotations were plotted.

**Extended Data Fig. 3.**
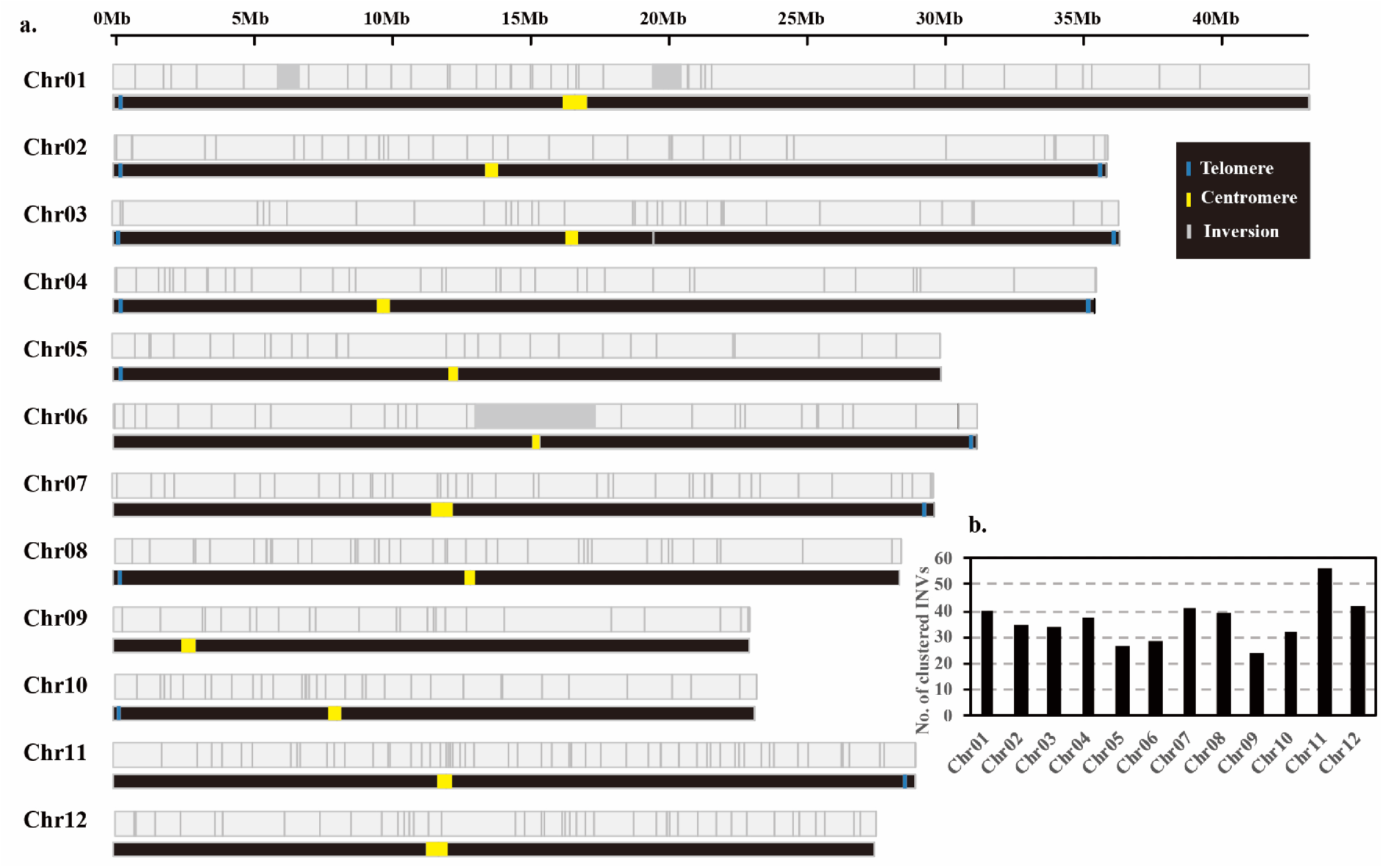
Chromosome distribution and amount of inversions. a. Genome-wide distribution of inversions based on the *GJ*-temp: IRGSP-1.0 genome. b. Number of inversions on 12 chromosomes based on the *GJ*-temp: IRGSP-1.0 genome.

**Extended Data Fig. 4.**
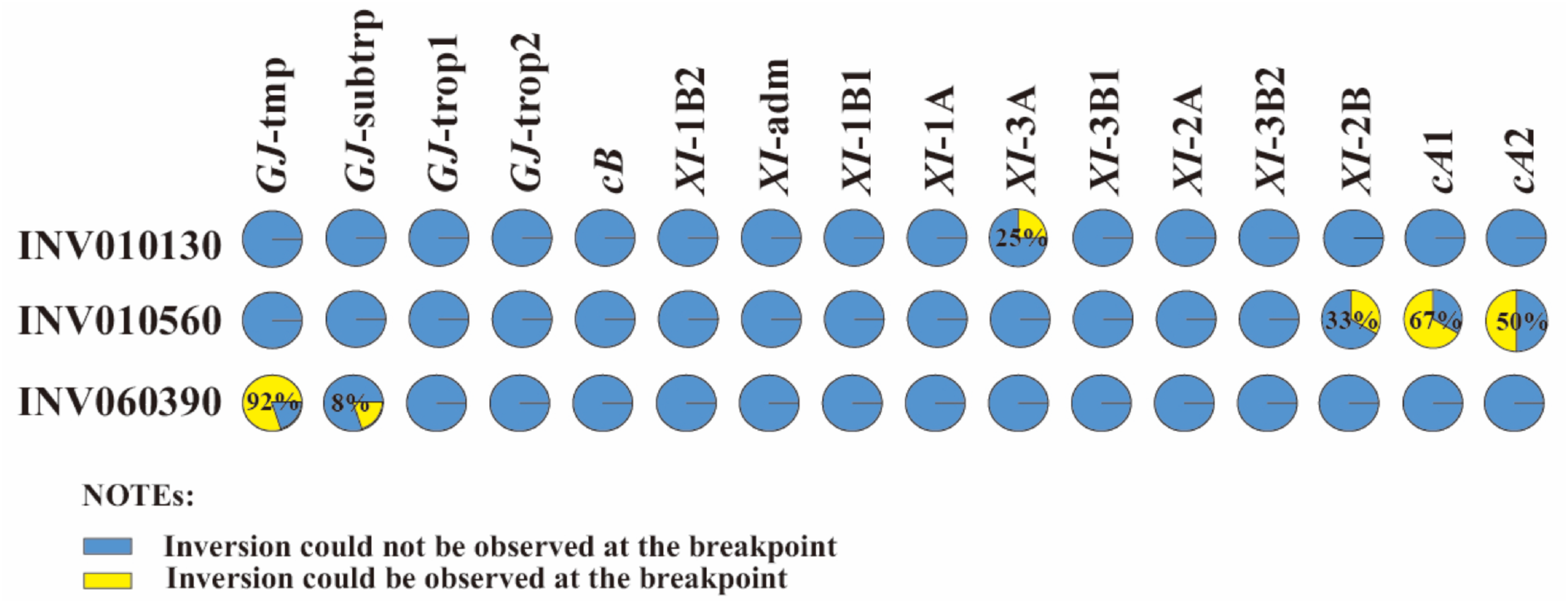
Validation of the 3 large (> 1 Mb) inversions in Asian rice, (INV010130, INV0101306 and INV060390) using NGS read data having at least 20× coverage from 192 3K-RGP samples.

**Extended Data Fig. 5.**
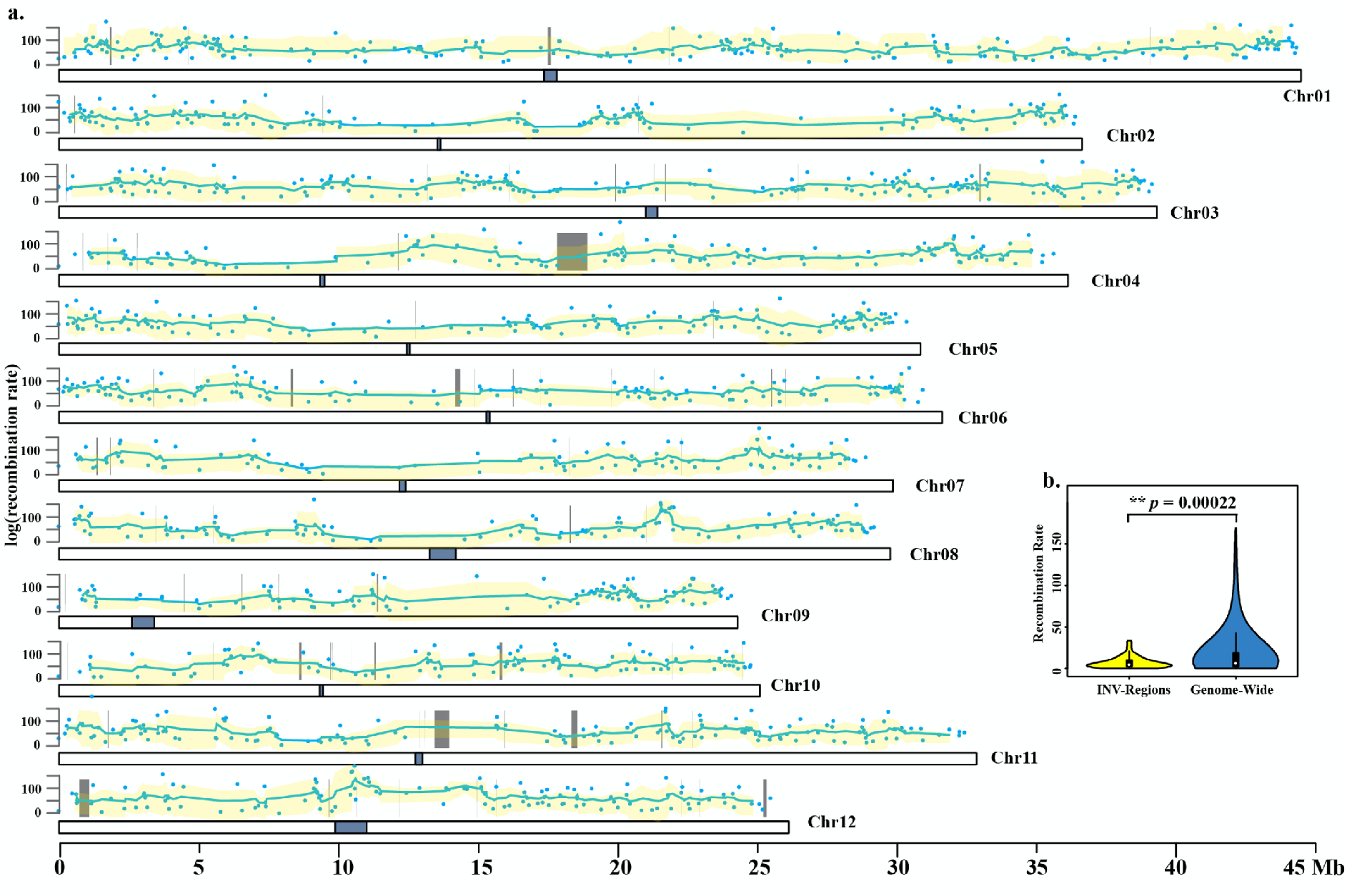
Recombination rate variation and inversion distribution in two *O. sativa* PSRefSeqs (*XI*-adm: MH63RS2 and *XI*-1A: ZS97RS2). a. Recombination rate of a RIL mapping population (Minghui 63 × Zhenshan 97). Dot points indicate the recombination rate and gray boxes indicate inversions. b. Comparison of recombination rate across inversions vs. genome-wide.

**Extended Data Fig. 6.**
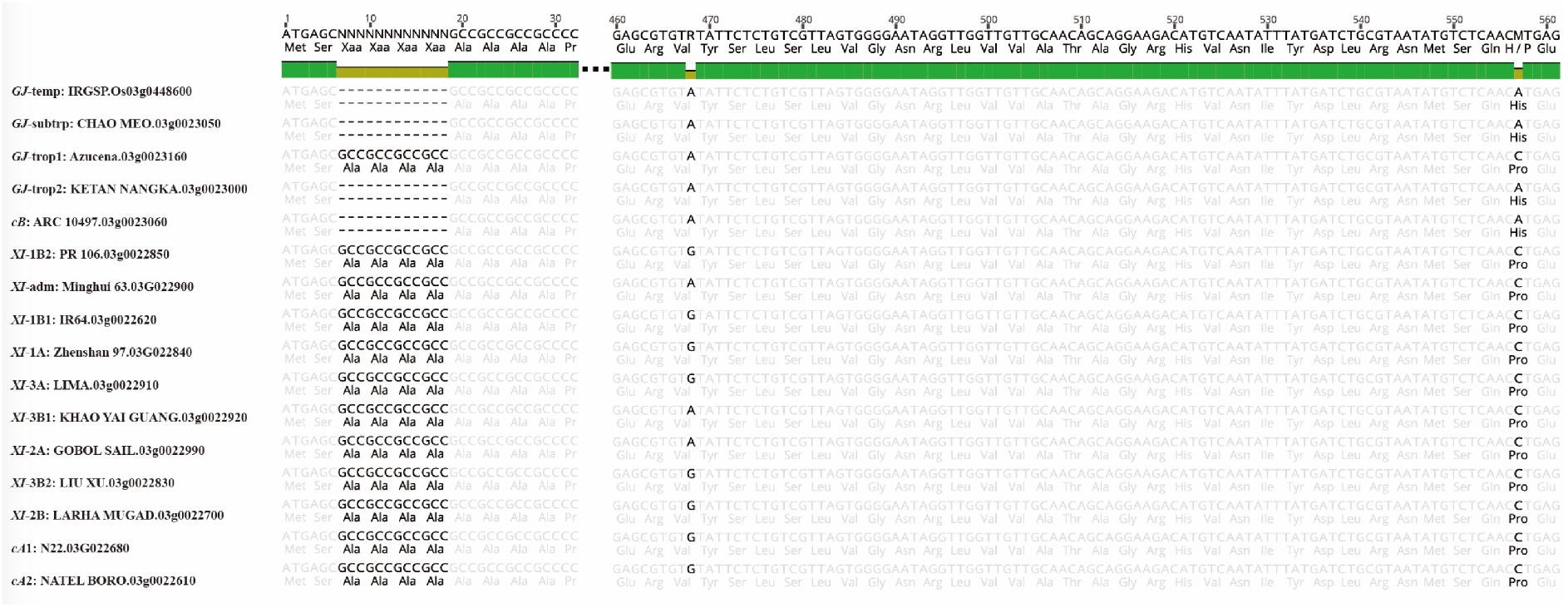
Two SNPs and a 12 bp InDel variant were observed within WD40-76 gene.

## Tables

**Table 1**. Assembly and annotation statistics of 18 *Oryza* reference genomes (18-genome data package).

**Table 2**. Summary of inversions identified across the 18-genome dataset.

## Supplemental Tables

**Supplementary Table 1**. Sequencing, data statistics of genome features, and BUSCO evaluation of *de novo* assemblies for 2 new wild *Oryza* genomes, *i*.*e. O. rufipogon* [AA] and *O. punctata* [BB].

**Supplementary Table 2**. Genome annotation statistics of 13 genomes. **Supplementary Table 3**. BUSCO assessments of genome annotations using both (a) transcriptome, and (b) protein model evidence.

**Supplementary Table 4**. Amount of PacBio Iso-Seq and RNA-Seq transcripts used for genome annotation.

**Supplementary Table 5**. Pan-genome inversions across the 18-genome data package, by comparing 15 *O. sativa* accessions, and 2 close relative genomes to the IRGSP-1.0. RefSeq.

**Supplementary Table 6**. Kolmogorov-Smirnov (KS) tests for genome-wide inversion distribution.

**Supplementary Table 7**. Asian rice subpopulation validation for specific inversions.

**Supplementary Table 8**. A summary of subpopulation specific and group specific inversions.

**Supplementary Table 9**. Details of 5 large inversions (> 1 Mb).

**Supplementary Table 10**. Summary of TE content of fine-scaled inverted regions based on 16 Asian rice genomes.

**Supplementary Table 11**. TE annotation of 16 *Oryza sativa* genomes.

**Supplementary Table 12**. 17 TEs from 4 superfamilies were observed with a higher amount (> 10 in this study) at inversion breakpoints of 16 Asian genomes.

**Supplementary Table 13**. Gene content analysis of inversions across 16 Asian rice genomes.

**Supplementary Table 14**. Comparison of transcript abundance levels for genes that were located within inversions, or at inversion breakpoints.

**Supplementary Table 15**. Seventy-eight inversions identified between *XI*-adm: MH63RS2 and *XI*-1A: ZS97RS2 genomes, the parents of a RIL-10 population. Note: To identify recombination rates, we only focused on inversions > 1 kb.

**Supplementary Table 16**. Cluster for inversions larger than 100 kb.

**Supplementary Table 17**. Investigation of 10 phenotypes. FT: flowering time from sowing. GWE: 100 grain weight (g). PL: panicle length (cm). PS: panicle shattering. SF: spikelet fertility. LS: leaf senescence. LW: leaf width. LL: leaf length. SIEC12: Salt injury at EC12. SIEC18: Salt injury at EC18.

**Supplementary Table 18**. Transcript abundance (FPKM value) of 8 genes located within ClusterINV92 (INV030400/INV030410/INV030420).

**Supplementary Table 19**. Validation of inversion ClusterINV92 and variants (3 SNPs and one 12 bp InDel) among different subpopulations by using 192 deep re-sequenced samples (> 20×).

**Supplementary Table 20**. Validation of 2,042 predicted inversions, of the same accessions used to create the Asian rice pan-genome, collected from the 3K-RGP data set.

**Supplementary Table 21**. A list of 6 inversions that were identified from the 3K-RGP data set, but were missed in the Asian rice pan-genome inversion index. INV3KSNPSEEK71124 and INV3KSNPSEEK71127, and INV3KSNPSEEK117247 and INV3KSNPSEEK117248 were found to be overlapping, and thus, were considered as a single inversion, resulting in 4 inversions were undetected in the pan-genome inversion index.

## References

1. Hossain, M. & Fischer, K. Rice research for food security and sustainable agricultural development in Asia: achievements and future challenges. GeoJournal 35, 286–298 (1995).

2. Wing, R.A., Purugganan, M.D. & Zhang, Q. The rice genome revolution: from an ancient grain to Green Super Rice. Nat Rev Genet 19, 505–517 (2018).

3. Vollset, S.E. et al. Fertility, mortality, migration, and population scenarios for 195 countries and territories from 2017 to 2100: a forecasting analysis for the Global Burden of Disease Study. The Lancet 396, 1285–1306 (2020).

4. Kirkpatrick, M. How and why chromosome inversions evolve. PLoS Biol 8, e1000501 (2010).

5. Wellenreuther, M. & Bernatchez, L. Eco-evolutionary genomics of chromosomal inversions. Trends in ecology evolution 33, 427–440 (2018).

6. Hoffmann, A.A. & Rieseberg, L.H. Revisiting the impact of inversions in evolution: from population genetic markers to drivers of adaptive shifts and speciation? Annual review of ecology, evolution, systematics 39, 21–42 (2008).

7. Sturtevant, A.H. A case of rearrangement of genes in Drosophila. Proc. Natl. Acad. Sci. USA 7, 235 (1921).

8. Dobzhansky, T. & Sturtevant, A.H. Inversions in the chromosomes of Drosophila pseudoobscura. Genetics 23, 28 (1938).

9. Barb, J.G. et al. Chromosomal evolution and patterns of introgression in Helianthus. Genetics 197, 969–979 (2014).

10. Volkert, F.C. & Broach, J.R. Site-specific recombination promotes plasmid amplification in yeast. Cell 46, 541–550 (1986).

11. Johnson, R.C. Bacterial site-specific DNA inversion systems. in Mobile DNA II 230–271 (American Society of Microbiology, 2002).

12. Hammer, M.F., Schimenti, J. & Silver, L.M. Evolution of mouse chromosome 17 and the origin of inversions associated with t haplotypes. Proc. Natl. Acad. Sci. USA 86, 3261–3265 (1989).

13. Flores, M. et al. Recurrent DNA inversion rearrangements in the human genome. Proc. Natl. Acad. Sci. USA 104, 6099–6106 (2007).

14. Hellen, E.H. Inversions and evolution of the human genome. eLS, 1-6 (2015).

15. Coughlan, J.M. & Willis, J.H. Dissecting the role of a large chromosomal inversion in life history divergence throughout the Mimulus guttatus species complex. Molecular ecology 28, 1343–1357 (2019).

16. Hey, J. Speciation and inversions: chimps and humans. Bioessays 25, 825–8 (2003).

17. Levy-Sakin, M. et al. Genome maps across 26 human populations reveal population-specific patterns of structural variation. Nat Commun 10, 1–14 (2019).

18. Wang, W. et al. Genomic variation in 3,010 diverse accessions of Asian cultivated rice. Nature 557, 43–49 (2018).

19. Rgp, K. The 3,000 rice genomes project. Gigascience 3, 7 (2014).

20. Fuentes, R.R. et al. Structural variants in 3000 rice genomes. Genome Res 29, 870–880 (2019).

21. Kou, Y. et al. Evolutionary genomics of structural variation in Asian rice (Oryza sativa) domestication. Molecular biology evolution 37, 3507–3524 (2020).

22. Stein, J.C. et al. Genomes of 13 domesticated and wild rice relatives highlight genetic conservation, turnover and innovation across the genus Oryza. Nat Genet 50, 285–296 (2018).

23. Zhang, J. et al. Extensive sequence divergence between the reference genomes of two elite indica rice varieties Zhenshan 97 and Minghui 63. Proc. Natl. Acad. Sci. USA 113, E5163–71 (2016).

24. Zhou, Y. et al. A platinum standard pan-genome resource that represents the population structure of Asian rice. Sci Data 7, 113 (2020).

25. Kawahara, Y. et al. Improvement of the Oryza sativa Nipponbare reference genome using next generation sequence and optical map data. Rice (N Y) 6, 4 (2013).

26. Huang, J. et al. Identifying a large number of high-yield genes in rice by pedigree analysis, whole-genome sequencing, and CRISPR-Cas9 gene knockout. Proc Natl Acad Sci U S A 115, E7559–E7567 (2018).

27. Xie, F. & Zhang, J. Shanyou 63: an elite mega rice hybrid in China. Rice (N Y) 11, 17 (2018).

28. Kou, Y. et al. Evolutionary genomics of structural variation in Asian rice (Oryza sativa) and its wild progenitor (O. rufipogon). BioRxiv (2019).

29. Kim, H. et al. Comparative physical mapping between Oryza sativa (AA genome type) and O. punctata (BB genome type). Genetics 176, 379–90 (2007).

30. Qin, P. et al. Pan-genome analysis of 33 genetically diverse rice accessions reveals hidden genomic variations. Cell 184, 3542–3558 (2021).

31. Longbiao, G. et al. Genetic Analysis and Utilization of the Important Agronomic Traits on Zhenshan 97* Minghui 63 Recombinant Inbred Lines (RIL) in Rice (Oryza sativa L.). Zuo wu xue bao 28, 644–649 (2002).

32. Huang, K. & Rieseberg, L.H. Frequency, Origins, and Evolutionary Role of Chromosomal Inversions in Plants. Frontiers in Plant Science 11, 296 (2020).

33. Li, C. et al. OsLBD37 and OsLBD38, two class II type LBD proteins, are involved in the regulation of heading date by controlling the expression of Ehd1 in rice. Biochemical Biophysical Research Communications 486, 720–725 (2017).

34. Žárský, V., Sekereš, J., Kubátová, Z., Pečenková, T. & Cvrčková, F. Three subfamilies of exocyst EXO70 family subunits in land plants: Early divergence and ongoing functional specialization. Journal of experimental botany 71, 49–62 (2020).

35. Ouyang, Y., Huang, X., Lu, Z., Yao, J. Genomic survey, expression profile and co-expression network analysis of OsWD40 family in rice. BMC Genomics 13, 100 (2012).

36. Itoh, J.I., Kitano, H., Matsuoka, M. & Nagato, Y. Shoot organization genes regulate shoot apical meristem organization and the pattern of leaf primordium initiation in rice. The Plant Cell 12, 2161–74 (2000).

37. McClintock, B. Cytological observations of deficiencies involving known genes, translocations and an inversion in Zea mays, (University of Missouri, College of Agriculture, Agricultural Experiment Station, 1931).

38. Ahn, S. & Tanksley, S. Comparative linkage maps of the rice and maize genomes. Proc. Natl. Acad. Sci. USA 90, 7980–7984 (1993).

39. Burke, J.M. et al. Comparative mapping and rapid karyotypic evolution in the genus helianthus. Genetics 167, 449–57 (2004).

40. Husband, B. Chromosomal variation in plant evolution. (JSTOR, 2004).

41. Levin, D.A. & Donald, A. The role of chromosomal change in plant evolution, (Oxford University Press, USA, 2002).

42. Crow, T. et al. Gene regulatory effects of a large chromosomal inversion in highland maize. PLoS Genet 16, e1009213 (2020).

43. Gage, J.L., Monier, B., Giri, A. & Buckler, E.S. Ten years of the maize nested association mapping population: impact, limitations, and future directions. The Plant Cell 32, 2083–2093 (2020).

44. The Computational Pan-Genomics Consortium. Computational pan-genomics: status, promises and challenges. Brief Bioinform 19, 118–135 (2018).

45. Vernikos, G.S. A Review of Pangenome Tools and Recent Studies. in The Pangenome: Diversity, Dynamics and Evolution of Genomes (eds. Tettelin, H. & Medini, D.) 89–112 (Cham (CH), 2020).

46. Athiyannan, N. et al. Long-read genome sequencing of bread wheat facilitates disease resistance gene cloning. Nat Genet 54, 227–231 (2022).

47. Bathke, J. & Lühken, G. OVarFlow: a resource optimized GATK 4 based Open source Variant calling workFlow. BMC bioinformatics 22, 1–18 (2021).

