## Supplementary information for "Pan-genome inversion index reveals evolutionary insights into the subpopulation structure of Asian rice (*Oryza sativa*)"

### Online Methods

#### The 18-genome Data Package

**Sequence and Assembly:** Seed and/or tissue from the 16 *O. sativa* (cultivated Asian rice), and 2 relatives (*O. rufipogon* and *O. punctata*) accessions were obtained from International Rice Research Institute (IRRI), Philippines, Huazhong Agricultural University (HZAU), China, and the International Center for Tropical Agriculture (CIAT), Colombia, and University of Arizona (UA), USA.

Genome assemblies for the 16 *O. sativa* accession were published previously[1-3]. The *O. rufipogon*, and updated *O. punctata* (BB genome) assemblies are reported here. The *O. rufipogon* accession (IRGC 106523) was selected as a "true" *O. rufipogon* representative based on its origin (Papua New Guinea, reproductively isolated from rice cultivation) and phenotype. A draft assembly of the BB genome species *O. punctata* (IRGC 105690) was previously reported[4] and upgraded *de novo* to a platinum standard reference sequence (PSRefSeq) here.

Briefly, the two genomes were sequenced to a minimum of 100 × coverage using PacBio long-read technology, assembled and validated to a PSRefSeq quality level following the identical strategy used for the 12 *O. sativa* genome data set previously described[3]. The Benchmarking Universal Single-Copy Orthologs (BUSCO v4.0) software package[5] was employed to evaluate the gene space completeness of each assembly.

**Genome Annotation:** Genome annotation used PacBio Iso-Seq and Illumina RNA-Seq data derived from RNA isolated root, panicle and young leaf tissue from Nipponbare and 13 newly sequenced *O. sativa* accessions[3] as "baseline" transcript evidence (*i.e.* RNA-Seq dataset#1). Annotation of Minghui 63 and Zhenshan 97 was reported previously with similar tissues[2]. In addition, we collected deep RNA-Seq from *O. sativa* cv. Nipponbare, Minghui 63 and Zhenshan 97 (RNA-Seq dataset#2[6]) for downstream transcriptome analyses. PacBio Iso-Seq data was deposited in NCBI under BioProject PRJNA760839. The RNA-Seq dataset#1 was deposited in NCBI under BioProject PRJNA659864. RNA-Seq dataset#2 were retrieved from NCBI BioProject PRJNA597070[6].

Protein coding genes for the 14 *O. sativa* genomes above were predicted using MAKER-P (3.01.03)[7] including expression evidence, homology and a*b initio* gene predictors FGENESH (v.8.0.0)[8,9], SNAP (0.15.7)[8,9]. Repeat masking was performed with RepeatMasker (http://www.repeatmasker.org) using a *Oryza* specific repeat library[4]. Expression evidence included reference guided transcript assemblies generated using StringTie (v1.3.4a)[10] and Cufflinks (v2.2.1)[10,11]. To generate assembled transcripts, quality inspected RNA-Seq reads from each library were mapped to their respective genomes using STAR (v2.5.3a)[12] with an iterative 2-pass mapping approach in which splice junctions generated from the first round were used to refine alignments in the subsequent round. Mapped reads from each library were merged, sorted, and indexed using SAMTools (v1.9)[13] to generate input for transcript assembly programs. All software packages were run with default options. High quality full length transcripts were clustered using Cd-hit (v4.6)[14] for 95% sequence identity using parameters -c 0.95 -n 10 -d 0 -M 3000. The clustered transcripts were further filtered for intron retention events using SUPPA2[15]. Additional transcript and homology evidence was used as described[4] to run MAKER-P with keep_preds option set to 1. The gene structure of the predicted models was further improved using PASA (v2.4.1)[16] using full length cDNA and EST's downloaded from GenBank with the query "EST[Keyword] AND *Oryza sativa*[Organism]". Functional domain identification was completed with InterProScan (v5.38-76.0)[17]. TRaCE[18] was used to assign canonical transcripts based on domain coverage, protein length, and similarity to transcripts assembled by Stringtie. The quality of annotations was assessed with MAKER-P generated Annotation Edit Distance (AED) values[19] and BUSCO, respectively. Only transcripts with AED scores < 1 were retained. Finally, gene annotations were imported to Ensembl core databases, verified, and validated for translation using the Ensembl API[20]. All genome annotations are available at Gramene Pan *Oryza* database (https://oryza.gramene.org/).

**Transposable Elements (TEs) Annotation**

We re-annotated all 16 cultivated *O. sativa*, and *O. rufipogon* and *O. punctata* genomes using the output of the latest version of the EDTA (v1.7.4)[21] TE annotation pipeline. The entire output was loaded into RepeatMasker (v 4.0.8)[22], with the exception of predicted helitron elements that were skipped because of a high false positive rate.

**Identification of Genomic Inversions and Assessments**

To discover large inversions (> 100 bp), we firstly tested four different analysis workflows (https://gitlab.kaust.edu.sa/zhouy0e/sv-for-o.sativa) on two genomes, *i.e.*, *GJ*-temp: IRGSP-1.0 and *XI*-adm: MH63RS2 (the 2nd genome version of Minghui 63 accession).

Workflow 1: The MH63RS2 genome assembly was split into overlapping reads of 50 kb in length at 5 kb step intervals, resulting in ~10 × coverage. The reads were then mapped onto the IRGSP-1.0 genome sequence using the tool CoNvex Gap-cost alignMents for Long Reads (NGMLR, v0.2.7)[23]. Inversions were called with SVIM (v1.1.0)[24], retaining only inversions with a depth greater > 6 that passed the caller's filtration criteria.

Workflow 2: Steps are same as in workflow 1 except that Sniffles (v1.0.7)[23] was used to call inversions.

Workflow 3: The MH63RS2 genome assembly was aligned to IRGSP-1.0 using Minimap2[25]. This step was followed by filtration for identity greater than 90% and length longer than 100 bp. Inversions were called using SyRI (*i.e.* Synteny and Rearrangement Identifier, v1.4)[26].

Workflow 4: Steps are the same as in workflow3 except the alignment tool used, *i.e.* Nucmer, part of the MUMmer (v4)[27].

To assess the accuracy of the four workflows, we followed two different strategies (https://gitlab.kaust.edu.sa/zhouy0e/sv-for-o.sativa). For Strategy 1, we took advantage of the fact that the orientation of the reads used was known. We checked all putative inversions for orientation of read mapping, and if an orientation was opposite to that of the original reads then the inversion was retained, otherwise it was filtered out. For strategy 2, dot-plots of the syntenic regions including the putative inversions were generated to visually validate the inversions.

Out of all workflows, workflow4 was selected based on our validation criteria.

In addition, Bionano optical maps of 13 *O. sativa* genomes[3] were used to validate the 5 inversions > 1 Mb.

Subsequently, the sequences of the 17 genomes were aligned to the IRGSP RefSeq with the MUMmer[27], filtering the alignments for a minimum identity of 90%, and minimum length of 100 bp, and the coordinates were retrieved using the fuction "show-coords" of MUMmer[27]. Finally, inversions were called using the SyRI tool (v1.4)[26] with default parameters, which provided VCFs (v.3) with ID, start, end of reference and query genome coordinates that was leveraged for pan-genome comparisons downstream.

**Pan-genome Inversions Index of the 18-genome Data Package**

Seventeen inversion vcf files derived from 17 genomes compared to IRGSP RefSeq were generated. Briefly, vcf files were sorted according to chromosome ID, coordinates and strand, and were merged using SURVIVOR (v1.0.7)[28], run under default parameters (https://github.com/fritzsedlazeck/SURVIVOR/wiki), except for the maximum allowed distance of 10 bp. In this case, inversions having start and end coordinates no more different than 10 bp were collapsed and considered as single inversions.

**Genome-wide Distribution of Pan-genome Inversion Index**

The distribution of pan-genome inversion index of Asian rice was studied. Inversions were further merged across pairwise assembly comparisons into a single set of coordinates. Deviations from a uniform distribution were then estimated on a per chromosome basis. The Kolmogorov-Smirnov (KS) test was performed on inversion start cooridnates[29,30]. In parallel, we performed a window-based analysis to determine whether clusters of inversions could be identified. To control for sample size, for each chromosome we performed 10,000 simulations of uniformly distributed positions of the same number as the inversions reported in that chromosome. From this, we determined the sample-size-specific false discovery rate of the KS test as well as the *p*-value for different numbers of observed inversions within windows of 200 kb in length. Window specific *p*-values were adjusted using the Benjamini-Hochberg procedure[31,32] to reduce the false positive rate on a per-chromosome basis. In addition, a

Bonferroni correction was applied to the significance threshold of KS tests to account for their application across multiple chromosomes.

**Species and Subpopulation Specific Inversions**

The pan-genome inversion index provided a complete list of all the inversions identified in our 18-genome data package. We then sorted out species-specific, group-specific, genome-specific candidates and shared inversions. Species-specific candidate inversions were divided into *O. punctata* (BB genome), *O. rufipogon* and *O. sativa* specific inversions, which refers to inversions that only could be detected in the *O. punctata*, *O. rufipogon* or *O. sativa* genomes. Group-specific candidates refers to inversions that are shared in multiple *O. sativa* genomes, *e.g.* only observed either in *Geng/Janponica* (*GJ)* subgroup, *Xian/Indica* (*XI*) subgroup, *circum-Aus* (*cA*) subgroup, or *circum-Basmati* subgroup genomes, respectively. Genome-specific candidate inversions refers to ones that could only be observed in one of 16 *O. sativa* genomes. The remaining inversions were defined as shared inversions.

Based on *O. sativa* subgroup-specific and genome-specific candidate inversions, we further studied if they were subpopulation(s) specific in Asian rice. These inversions were further validated using 192 deep re-sequenced samples (> 20 ×) derived from the 3K-RGP[33] dataset selected from each subpopulation (12 samples per subpopulation). By studying the 2 kb-wide alignment patterns at breakpoints in these accessions, we collected the frequency of each single inversion in different subpopulations. Based on these observations, inversions were classified into four groups: 1) genome-specific inversions, *i.e.* the inversions could be observed in only one of the query genomes but not in any of the 192 accessions; 2) subpopulation specific inversions, *i.e.* the inversions could be observed in one of the query genomes and also in one single subpopulation; 3) near-subpopulation specific inversions, *i.e.* the inversions could be observed in one of the query genomes, and mainly observed in one single subpopulation, but also could be found in other subpopulations with a lower frequency, and 4) subpopulation shared inversions, *i.e.* the inversions could be observed in more than one subpopulation.

**Inversion Rate Estimation**

To estimate inversion rates, we used a pair of genomes with estimated divergence times to a most recent common ancestor (MRCA), and divided the total number of inversion by twice the time to the MRCA (corresponding to the total branch length of the genealogy on two nodes). We applied this method to obtain three estimates:

1) using BB vs AA genomes (*i.e. O. punctata* vs. *O. sativa GJ*-temp IRGSP), with an estimated divergence time of 2.5 million years ago (MYA), the estimate is: 316 / (2*2.5 MY) = 63.2 inversions per MY;

2) using domesticated and AA progenitor genomes (*i.e. O. sativa GJ*-temp IRGSP vs. *O. rufipogon*) with an estimated divergence time of 0.5 MYA, the estimate is: 194 / (2* 0.5 My) = 194 inversions per MY;

3) using two *GJ* domesticated AA genomes (*i.e. O. sativa GJ*-trop2 KETAN NANGKA vs. *O. sativa GJ*-temp IRGSP, which showed the largest genetic distance among the *GJ*- subgroup) with an estimated divergence time of 0.01 MYA, using non-shared inversions to avoid over estimation, and the estimate is: 22 / (2* 0.01 My) = 1,100 inversions per MY.

**Differential Transcript Abundance (DTAs) Analysis within Inversions**

To investigate the effect of an inversion on gene expression, we searched for differences in transcript abundance within inversions in Asian rice based on two RNA-Seq datasets (dataset#1 and dataset#2). FPKM (Fragments Per Kilobase of exon model per Million mapped fragments) values were identified following an accurate pipeline[34], which included HISAT2[35] for alignment, and StringTie (v1.3.4a)[10] to obtain the normalized values (FPKM). Taking into account different transcript abundances between the two RNA-Seq datasets (2 Gb for dataset#1 vs 6 Gb for dataset#2), we used different minimum filtration of, *i.e.,* FPKM > 0.1 in RNA-Seq dataset#1 and FPKM > 1 in RNA-Seq dataset#2. Differential Transcript Abundance (DTA) of dataset#2, was carried out by edgeR[36] with P value < 0.01 and abundance change > 2.

**Genome Recombination Rate of Inversions**

To study the genome recombination rate of inversions, we used genetic data based on a RIL population derived from accessions *O. sativa cv. XI*-adm: Minghui 63 and *XI*-1A: Zhenshan 97[37,38]. Briefly, re-seuqencing data was obtained from 210 RILs and a

Bin Map containing 1,619 bins was generated. The physical positions of each bin were derived from updated versions both the MH63RS2 and ZS97RS2 reference genomes[38]. Then, a genetic map based on the RILs panel of 1,619 bins (Supplementary Note Table 1) was constructed using the MSTMap algorithm[39]. By comparing genetic and physical distances between neighboring Bin Map markers, we estimated the relative changes of the recombination rate of the two genomes[40]. Then, we compared recombination rates in the bins that overlapped with inversions and genome-wide, respectively.

#### Supplementary Notes

##### Supplementary Note 1: The 18-genome Data Package: Sequencing, Assembly, Annotation and Visualization

##### Sequence and Assemblies

The pan-genome of Asian rice includes 16 ultra high-quality genomes that represent the sub-population structure level of the 3K-RGP data set, including: 1) 15 platinum stranded reference sequences (PSRefSeqs), and 2) the previously published IRGSP RefSeq[1-3] (Table 1).

In addition, chromosome-level reference assemblies for two wild species - *i.e. O. rufipogon* [AA], and *O. punctata* [BB] were generated using long-read sequencing technology. Both genomes were sequenced with > 100 × long-read genome coverage using the PacBio Sequel II platform. These assemblies have sizes of 461 and 422 Mb, with contig N50s of 32.5 Mb and 33.1 Mb, respectively (Supplementary Table 1). Only 7 and 16 gaps remain for the *O. rufipogon* and *O. punctata* genomes (Supplementary Table 1), respectively. Data for these new genomes can be found in Genbank (https://www.ncbi.nlm.nih.gov/) under public BioProjects PRJNA609053 (*O. rufipogon*) and PRJNA13770 (*O. punctata*) (Supplementary Table 1).

BUSCO scores for the two new assemblies were 97.90% (*O. rufipogon*) and 97.00% (*O. punctata*) (Supplementary Table 1, Supplementary Note Table 2). Of note, 16 BUSCO genes were missing in the *O. rufipogon* genome sequence, which were also absent in the all *O. sativa* genomes as well[3]. However, 2 (EOG093605AK and EOG09360AWY) out of these 16 genes could be identified in *O. punctata* (BB genome) (Supplementary Note Table 2). Combined with our previous analysis with Asian rice and maize[3], 14 BUSCO genes are absent in the *Oryza* genus, 12 of which are absent in both the *Oryza* genus and *Zea mays* (Supplementary Note Table 2).

The assembly and conserved gene content statistics for the *O. rufipogon* and *O. punctata* genomes demonstrate their high-quality, genome-wide contiguity, and completeness. Hence, we regard both assemblies as PSRefSeqs as previously reported in Asian rice[1-3].

#### Asian Rice Pan-genome Annotation

To minimize bias associated with different methods of gene and repeat finding, we applied a uniform set of annotation protocols (implemented through MAKER-P[7]) to 14 Asian rice PSRefSeqs (see methods). To accomplish this task integrated both Illumina baseline RNA-Seq data (RNA-seq database#1) with PacBio Iso-Seq data, isolated independently from similar tissues (*i.e.* young leaves, roots and panicles) (Supplementary Table 4). As a result, we annotated on average 36,347 genes, with an average length 3,448 bp (Supplementary Table 2). These annotations resulted in an expected AED distribution (Extended Data Fig. 2), and BUSCO scores greater than 97.5% for both transcript and protein models (Supplementary Table 3).

#### 18-Genome Data Package Visualization

For alignment, analysis, visualization, and public availability, we uploaded the 18-genome data package into the Persephone® multi-genome browser (https://web.persephonesoft.com/). The types of data tracks visualized include gene models, marker locations, BLAST matches, RNA-Seq coverage and sequence tracks. As the genomes are closely related, the maps could be aligned by connecting short sequence tags (100 bp) derived from the IRGSP-1.0 RefSeq and mapped onto the other 17 genomes using BLASTN. An example alignment of 18 maps from chromosome 1, and an 800 kb region that includes inversion cluster 92 on chromosome 3 are shown in Extended Data Fig. 1.

#### Transposable Element Content

We re-annotated transposable elements (TEs) and investigated TE related sequence (TE-RS) content in the inversions across our 18-genome data set. The overall amount of TE-RSs ranged from 47.41% (*O. sativa XI*-3A LIMA) to 56.54% (*O. rufipogon*) with an average of 51.26% across all 18 genomes (Table 1, Supplementary table 11).

**Supplementary Note 2: Genomic Inversion Identification Workflow**

To discover inversions, we first tested four different analysis workflows (see methods) by comparing two *O. sativa* genomes, *i.e., GJ*-temp: IRGSP-1.0 and *XI*-adm: MH63RS2 genomes. We identified 25, 131, 55 and 235 raw inversions based on the 4 workflows, respectively (Supplementary Note Table 3). Following assessments by dotplots and long sequence (50 Kb) mapping, 22 (72%), 39 (29.8%), 33 (60%) and 178 (75.7%) raw inversions were confirmed (Supplementary Note Fig. 1a) for 4 workflows, respectively, and the false positives ones were filtered out (Supplementary Note Fig. 1b). In comparison to workflow 4, workflows 1, 2 and 3 identified only 10%, 21.7% and 18.3% of all inversions, respectively (Supplementary Note Table 3). Upon inspection, we determined that workflow 4 captured the overall majority of true large inversions.

**Supplementary Note 3: Chromosomal Distribution of Pan-genome Inversions Index**

To study the chromosomal distribution of pan-genome inversions, we further clustered all overlapping inversions located less than 15 kb apart. In doing so, we obtained 435 clustered inversions, ranging from a minimum of 27 (Chr09) to a maximum of 56 (Chr11) (SD = 8.15) (Extended Data Fig. 3). We tested for uniformity of inversion distribution across the 12 rice chromosomes using the Kolmogorov-Smirnov (KS) test. No significant deviation from uniformity was found in any chromosome. Window-based (200 kb) analyses also failed to reveal any significant clustering of inversion positions (Supplementary Table 6). These results demonstrate that the large inversions (> 100 bp) detected in this study are evenly distributed across each genome studied.

**Supplementary Note 4: The 5 Largest Inversions**

**Identification and Description**

We identified five inversions greater than 1 Mb detected relative to the IRGSP RefSeq (*i.e.* INV010130, INV010560, INV060390, INV080710 and INV100690). These five inversions span a total of 10.80 Mb and comprise 992 genes (2.96 Mb, 27.50% of total inverted region) and 11,956 transposable element related fragments (6.25 Mb, 61% of total inverted region) (Supplementary table 9).

Taking advantage of the Bionano optical maps from 13 *O. sativa* genomes, three *O. sativa* inversions (INV010130, INV010560 and INV060390) could be independently validated (Fig. 2). INV010130 (Chr01:5,966,062-7,971,800, length: ~2.0 Mb) could only be identified in a single genome (*i.e. XI*-3A: LIMA) relative to all remaining 17 assemblies (Supplementary Table 8-9, Fig. 2a). INV010560 (Chr01:19,913,222-21,736,006, length: ~1.8 Mb) could be identified in two genomes (*cA*1: N22 and *cA*2: NATEL BORO) relative to all remaining 16 genomes (Supplementary Table 5 & 9, Fig. 2b). Finally, INV060390 (Chr06: 13,119,682-17,632,495, length: 4.5 Mb) was detected in all 17 genomes relative to the IRGSP RefSeq, which means that this inversion arose in the *GJ*-temp: IRGSP RefSeq only (Supplementary Table 5 & 9, Fig. 1c). This inversion was first reported by a comparison of the *GJ*-temp: IRGSP RefSeq with the R498 "indica" genome sequence[41].

Two other inversions (INV080710 and INV100690) were detected in only *O. rufipogon* and *O. punctata*, respectively (Supplementary Table 5 & 9). INV080710 (Chr08: 19,521,753-20,651,007, length: 1.13 Mb) could only be identified in *O. punctata*, which was first reported by physical mapping and also confirmed by fluorescence in situ hybridization (FISH) analyis[42]. INV100690 (Chr10: 16,380,643-17,708,643, length: 1.33 Mb), INV010130 and INV010560 are reported for the first time in this study.

To determine the possible effects of these inversions on transcript abundance, we investigated genes located at or near all inversion breakpoints and detected a single breakpoint associated with INV010560 (Chr01: 19,913,222-21,736,006, length: 1.8 Mb) that was found to lie within the gene- *Rpr2/Rpp21* (Os01g0541600, function unknown) (Supplementary Note Fig. 2). The inversion breakpoint is located in an intron between *Rpr2/Rpp21* wo exons, thereby translocating exon 1 and moving it ~1.4 Mb away from exon 2, thereby disrupting the coding sequence. Analysis of 'baseline' RNA-Seq data (Online methods, dataset#1, *i.e.* panicle, leaf and root) from the two varieties that contain this inversion (*O. sativa* cv. *cA*1: N22 and *cA*2: NATEL BORO) showed, as predicted, a complete absence of transcript evidence, whereas transcription was detected in all similar *O. sativa* cv. Nipponbare tissues (Supplementary Note Fig. 2). The phenotypic consequences of this gene ablation, if any, remains to be determined.

**Subpopulation Validation**

The three largest inversions from *O. sativa* were further validated using the 3K-RGP dataset[33]. We selected 192 samples (> 20 ×), which include 12 samples from each Asian rice subpopulation, and studied their alignment patterns at each breakpoint (2 kb). We found that INV010130, identified in only genome *XI*-3A: LIMA, can also be observed in 3 out of the 12 (25%) samples from *XI*-3A subpopulation (Extended Data Fig. 4). INV010560, identified in *cAus* genomes *cA*1: N22 and *cA*2: NATEL BORO, can be mainly identified in the respective subpopulations *cA*1 (50%, 6 out of 12 tested samples) and *cA*2 (67%, 8 out of 12 tested samples) subpopulation, but also was identified in the *XI*-2B subpopulation (4 out of 12 tested samples) (Extended Data Fig. 4). The largest inversion, INV060390 (~5 Mb), which was identified in genome *GJ*-temp: IRGSP-1.0, could be mainly observed in the *GJ*-temp subpopulation (91.67%, 11 out of 12 tested samples), and it is also found at a low frequency in the *GJ*-subtrp subpopulation (8.33%, 1 out of 12 tested samples) (Extended Data Fig. 4).

**Supplemental Note Figs**

**Supplementary Note Fig. 1**. Assessments of inversions across 18 PSRefSeqs. True positives show (a) clean inverted regions and breakpoints. False positives (b) are mostly due to the incorrect assessment of tandemly repeated regions.

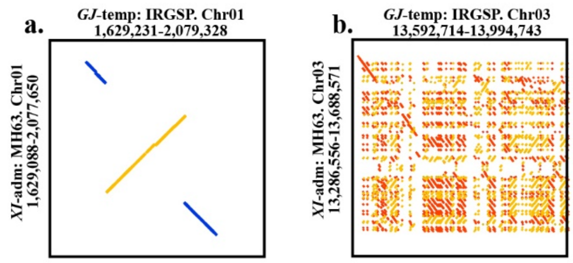


**Supplementary Note Fig. 2**. Disruption of gene expression by inversions.
a. A *Rpr2/Rpp21* gene (Os01g0541600) was observed at the breakpoint of INV010560 (1.82 Mb) and two exons were spited by the inversion.
b. A complete disruption of the *Rpr2/Rpp21* gene resulting in transcript ablation.

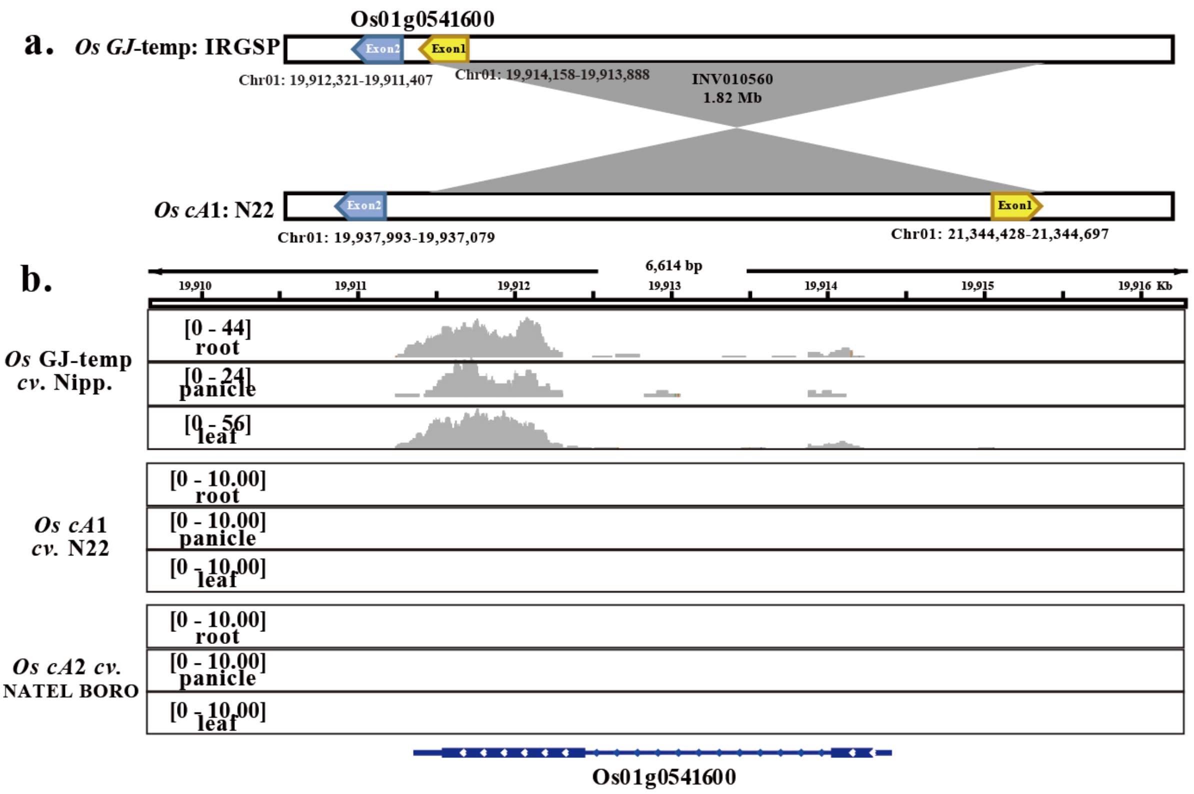

**Supplemental Note Tables**

**Supplementary Note Table 1**. The physical distance and genetic distance of bin map markers for the RIL population[43] that derived from Minghui 63 and Zhenshan 97.

**Supplementary Note Table 2**. BUSCO evaluation of the *O. rufipogon* [AA] and *O. punctata* [BB] genomes compared with Asian rice pan-genome missing genes[3].

**Supplementary Note Table 3**. The assessment of 4 workflows used to identify genomic inversions between the IRGSP-1.0 RefSeq and MH63V2 genome sequences.

#### References


1. Kawahara, Y. *et al.* Improvement of the Oryza sativa Nipponbare reference genome using next generation sequence and optical map data. *Rice (N Y)* **6**, 4 (2013).
2. Zhang, J. *et al.* Extensive sequence divergence between the reference genomes of two elite indica rice varieties Zhenshan 97 and Minghui 63. *Proc. Natl. Acad. Sci. USA* **113**, E5163-71 (2016).
3. Zhou, Y. *et al.* A platinum standard pan-genome resource that represents the population structure of Asian rice. *Sci Data* **7**, 113 (2020).
4. Stein, J.C. *et al.* Genomes of 13 domesticated and wild rice relatives highlight genetic conservation, turnover and innovation across the genus Oryza. *Nat Genet* **50**, 285-296 (2018).
5. Simao, F.A., Waterhouse, R.M., Ioannidis, P., Kriventseva, E.V. & Zdobnov, E.M. BUSCO: assessing genome assembly and annotation completeness with single-copy orthologs. *Bioinformatics* **31**, 3210-2 (2015).
6. Zhao, L. *et al.* Integrative analysis of reference epigenomes in 20 rice varieties. *Nat Commun* **11**, 2658 (2020).
7. Campbell, M.S., Holt, C., Moore, B. & Yandell, M. Genome annotation and curation using MAKER and MAKER‐P. *Current protocols in bioinformatics* **48**, 4.11. 1-4.11. 39 (2014).
8. Salamov, A.A. & Solovyev, V.V. Ab initio gene finding in Drosophila genomic DNA. *Genome Res* **10**, 516-22 (2000).
9. Korf, I. Gene finding in novel genomes. *BMC Bioinformatics* **5**, 59 (2004).
10. Pertea, M. *et al.* StringTie enables improved reconstruction of a transcriptome from RNA-seq reads. *Nat Biotechnol* **33**, 290-5 (2015).
11. Trapnell, C. *et al.* Differential gene and transcript expression analysis of RNA-seq experiments with TopHat and Cufflinks. *Nat Protoc* **7**, 562-78 (2012).
12. Dobin, A. *et al.* STAR: ultrafast universal RNA-seq aligner. *Bioinformatics* **29**, 15-21 (2013).
13. Li, H. *et al.* The Sequence Alignment/Map format and SAMtools. *Bioinformatics* **25**, 2078-9 (2009).
14. Li, W. & Godzik, A. Cd-hit: a fast program for clustering and comparing large sets of protein or nucleotide sequences. *Bioinformatics* **22**, 1658-9 (2006).
15. Trincado, J.L. *et al.* SUPPA2: fast, accurate, and uncertainty-aware differential splicing analysis across multiple conditions. *Genome Biol* **19**, 40 (2018).
16. Haas, B.J. *et al.* Improving the Arabidopsis genome annotation using maximal transcript alignment assemblies. *Nucleic acids research* **31**, 5654-5666 (2003).
17. Jones, P. *et al.* InterProScan 5: genome-scale protein function classification. *Bioinformatics* **30**, 1236-40 (2014).
18. Olson, A.J. & Ware, D. Ranked Choice Voting for Representative Transcripts with TRaCE. *Bioinformatics* (2021).
19. Campbell, M.S. *et al.* MAKER-P: a tool kit for the rapid creation, management, and quality control of plant genome annotations. *Plant Physiol* **164**, 513-24 (2014).
20. Stabenau, A. *et al.* The Ensembl core software libraries. *Genome Res* **14**, 929-33 (2004).
21. Ou, S. *et al.* Benchmarking transposable element annotation methods for creation of a streamlined, comprehensive pipeline. *Genome Biol* **20**, 275 (2019).
22. Tarailo-Graovac, M. & Chen, N. Using RepeatMasker to identify repetitive elements in genomic sequences. *Curr Protoc Bioinformatics* **4**, 4-10 (2009).

23. Sedlazeck, F.J. *et al.* Accurate detection of complex structural variations using single-molecule sequencing. *Nat Methods* **15**, 461-468 (2018).
24. Heller, D. & Vingron, M. SVIM: structural variant identification using mapped long reads. *Bioinformatics* **35**, 2907-2915 (2019).
25. Li, H. Minimap2: pairwise alignment for nucleotide sequences. *Bioinformatics* **34**, 3094-3100 (2018).
26. Goel, M., Sun, H., Jiao, W.B. & Schneeberger, K. SyRI: finding genomic rearrangements and local sequence differences from whole-genome assemblies. *Genome Biol* **20**, 277 (2019).
27. Delcher, A.L., Salzberg, S.L. & Phillippy, A.M. Using MUMmer to identify similar regions in large sequence sets. *Current protocols in bioinformatics* **00**, 10.3.1-10.3.18 (2003).
28. Jeffares, D.C. *et al.* Transient structural variations have strong effects on quantitative traits and reproductive isolation in fission yeast. *Nat Commun* **8**, 14061 (2017).
29. Massey Jr, F.J. The Kolmogorov-Smirnov test for goodness of fit. *Journal of the American statistical Association* **46**, 68-78 (1951).
30. Virtanen, P. *et al.* SciPy 1.0: fundamental algorithms for scientific computing in Python. *Nat Methods* **17**, 261-272 (2020).
31. Benjamini, Y. & Hochberg, Y. Controlling the false discovery rate: a practical and powerful approach to multiple testing. *Journal of the Royal statistical society: series B* **57**, 289-300 (1995).
32. Seabold, S. & Perktold, J. Statsmodels: Econometric and statistical modeling with python. in *Proceedings of the 9th Python in Science Conference* Vol. 57 61 (Austin, TX, 2010).
33. 3K-RGP. The 3,000 rice genomes project. *Gigascience* **3**, 7 (2014).
34. Pertea, M., Kim, D., Pertea, G.M., Leek, J.T. & Salzberg, S.L. Transcript-level expression analysis of RNA-seq experiments with HISAT, StringTie and Ballgown. *Nat Protoc* **11**, 1650-67 (2016).
35. Kim, D., Langmead, B. & Salzberg, S.J.N.m. HISAT: a fast spliced aligner with low memory requirements. *Nat Methods* **12**, 357-360 (2015).
36. Robinson, M.D., McCarthy, D.J. & Smyth, G.K. edgeR: a Bioconductor package for differential expression analysis of digital gene expression data. *Bioinformatics* **26**, 139-40 (2010).
37. Hua, J. *et al.* Genetic dissection of an elite rice hybrid revealed that heterozygotes are not always advantageous for performance. *Genetics* **162**, 1885-1895 (2002).
38. Zhou, G. *et al.* Genetic composition of yield heterosis in an elite rice hybrid. *Proc. Natl. Acad. Sci. USA* **109**, 15847-15852 (2012).
39. Wu, Y., Bhat, P.R., Close, T.J. & Lonardi, S. Efficient and accurate construction of genetic linkage maps from the minimum spanning tree of a graph. *PLoS Genet* **4**, e1000212 (2008).
40. Wu, J. *et al.* Physical maps and recombination frequency of six rice chromosomes. *The Plant Journal* **36**, 720-730 (2003).
41. Du, H. *et al.* Sequencing and de novo assembly of a near complete indica rice genome. *Nat Commun* **8**, 1-12 (2017).
42. Kim, H. *et al.* Comparative physical mapping between Oryza sativa (AA genome type) and O. punctata (BB genome type). *Genetics* **176**, 379-90 (2007).
43. Longbiao, G. *et al.* Genetic Analysis and Utilization of the Important Agronomic Traits on Zhenshan 97* Minghui 63 Recombinant Inbred Lines (RIL) in Rice (Oryza sativa L.). **28**, 644-649 (2002).
